# Democratizing Protein Language Model Training, Sharing and Collaboration

**DOI:** 10.1101/2024.05.24.595648

**Authors:** Jin Su, Zhikai Li, Tianli Tao, Chenchen Han, Yan He, Fengyuan Dai, Qingyan Yuan, Yuan Gao, Tong Si, Xuting Zhang, Yuyang Zhou, Junjie Shan, Xibin Zhou, Xing Chang, Shiyu Jiang, Dacheng Ma, The OPMC, Martin Steinegger, Sergey Ovchinnikov, Fajie Yuan

**Affiliations:** Zhejiang University, Hangzhou, China; School of Engineering, Westlake University, Hangzhou, China; Suzhou Polynovo Biotech Co., Ltd., Suzhou, China; Shenzhen Institute of Synthetic Biology, Shenzhen Institutes of Advanced Technology, Chinese Academy of Sciences, Shenzhen, China; Zhejiang Lab, Hangzhou, China; Open Protein Modeling Consortium; School of Biological Sciences, Seoul National University, Seoul, South Korea; Massachusetts Institute of Technology, Cambridge, USA

## Abstract

Training and deploying large-scale protein language models typically requires deep machine learning expertise — a barrier for researchers outside this field. SaprotHub overcomes this challenge by offering an intuitive platform that facilitates training and prediction as well as storage and sharing of models. We provide the ColabSaprot framework built on Google Colab which potentially powers hundreds of protein training and prediction applications, enabling researchers to collaboratively build and share customized models.

---

Proteins are fundamental to virtually all biological processes and central to medicine and biotechnology [1, 2, 3]. Despite this centrality, deciphering protein structure and function has remained a formidable challenge. This landscape was recently transformed by two breakthroughs: The success of AlphaFold2 [4] ushered in a new era for structural biology by predicting structures with experimental-level accuracy, while in parallel, large-scale protein language models (PLMs) are driving unprecedented advances in function prediction.

This progress is driven by a suite of powerful PLMs that have demonstrated remarkable efficacy across diverse tasks [5, 6, 7, 8, 9, 10, 11, 12, 13, 14]. However, leveraging these advanced models presents significant technical hurdles for researchers without extensive machine learning (ML) expertise. The challenges span the entire workflow, from model selection and data preprocessing to the training and evaluation of models with billions of parameters. This complexity creates a critical barrier, hindering broader adoption and innovation by the very researchers who stand to benefit most.

ColabFold [15] addressed a similar accessibility barrier for structure prediction by deploying AlphaFold2 on Google Colab, effectively democratizing its use. Despite this success, a critical gap remains: ColabFold does not support the more complex task of training custom models for function prediction.

To bridge this gap, we introduce ColabSaprot and SaprotHub, a platform designed specifically for protein function prediction. Built on Google Colab, ColabSaprot empowers researchers without ML expertise to train their own task-specific PLMs through an intuitive interface. Crucially, the platform supports a broad spectrum of prediction tasks, ensuring its utility extends far beyond single-task applications.

Complementing this user-friendly platform, we introduce the *Open Protein Modeling Consortium* (OPMC), an initiative to foster a collaborative ecosystem for community-driven protein language modeling (see Supplementary Information). The OPMC framework enables researchers to share their bespoke models, fine-tune existing ones contributed by peers, or apply them directly for their own research. This creates a virtuous cycle of sharing, refinement, and application, accelerating collective progress in the field. As the inaugural platform integrated with OPMC, SaprotHub represents the first step in realizing this vision for community-centric AI development.

This work comprises three key contributions: a foundation PLM named Saprot [14] (Fig. 1a,b), Colab-Saprot (Fig. 2a,b,c) enabling easy training (or fine-tuning) and inference of Saprot on the Colab platform through the adapter learning technique [16, 17], and SaprotHub, a community repository (Fig. 2d) for storing, sharing, searching, and collaborative development of fine-tuned Saprot models. By integrating advanced protein language models, cloud-based computing on Colab, and adapter-based fine-tuning techniques, it addresses several key challenges—namely, the difficulty of sharing and collectively utilizing large-scale PLMs, the risk of parameter catastrophic forgetting during continual learning [18], and the need to protect proprietary biological data when sharing results. Below, we detail these three modules.

**Fig. 1:**
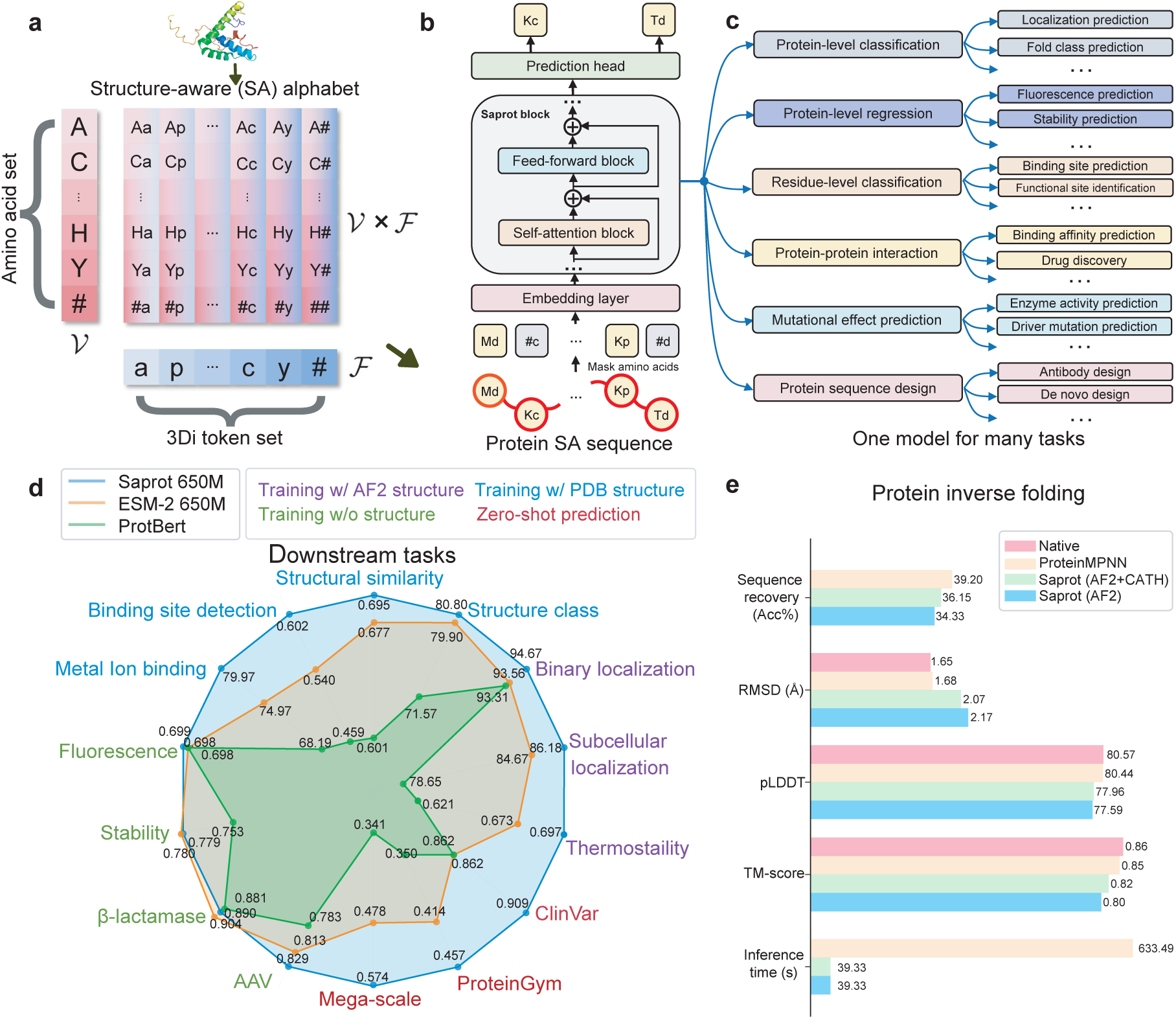
Illustration of Saprot. **a,** The proposed SA alphabet. #a, #p,…,#y represent only the 3Di token being visible, while A#, C#,…, Y# represent the AA token being visible. **b,** Model architecture of Saprot. **c,** Saprot supports a wide range of protein prediction tasks, see more tasks in Supplementary Table 1. **d,** Performance comparison on 14 diverse prediction tasks. **e.** Performance on protein sequence design. “AF2+CATH” indicates that Saprot was pre-trained using AlphaFold2 predicted structures and fine-tuned with experimental structures from CATH. “AF2” is a version of Saprot trained only with the AlphaFold2 predicted structure. CATH is also used for training ProteinMPNN.

**Fig. 2:**
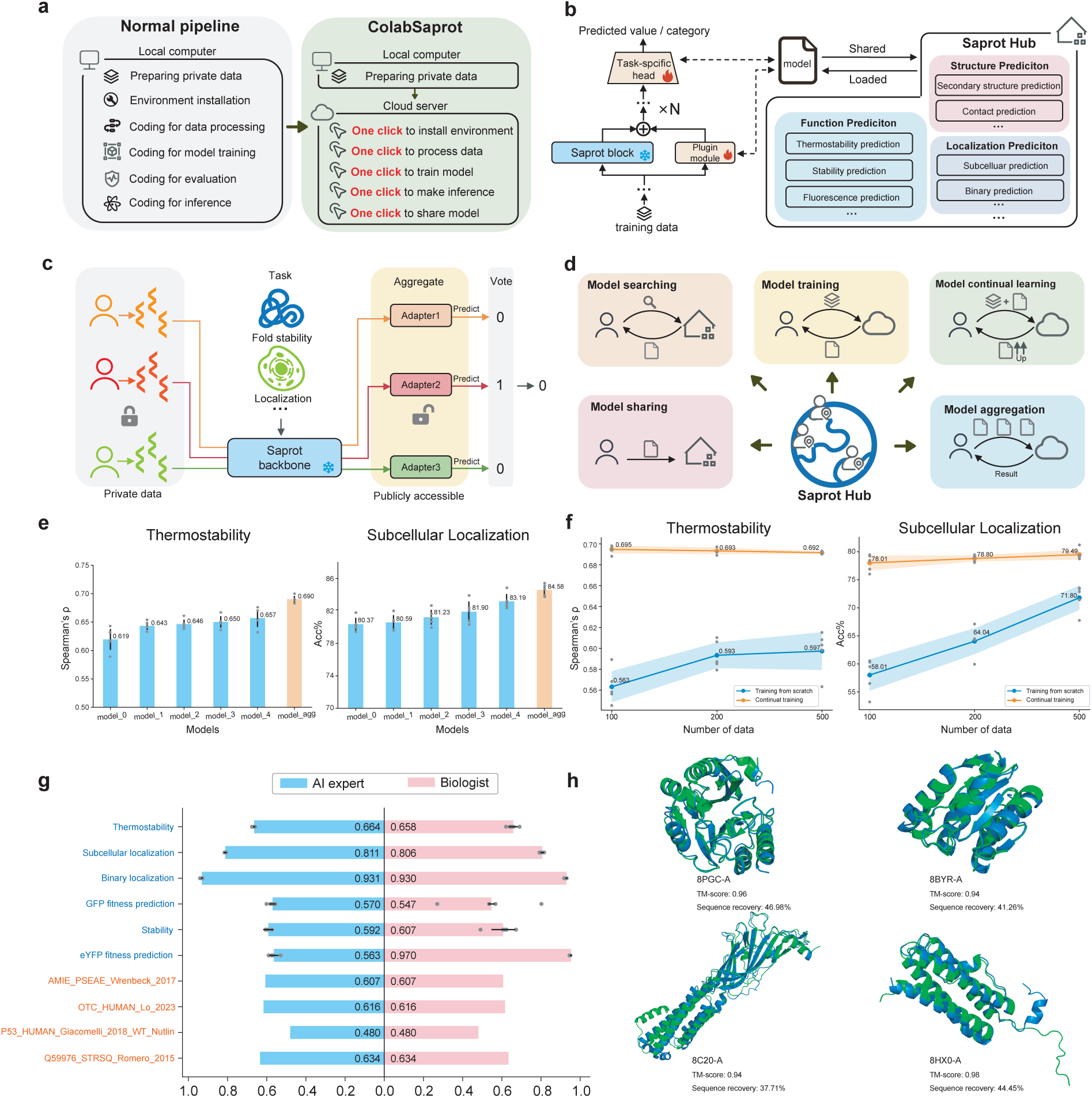
Illustration of ColabSaprot and SaprotHub. **a,** Comparison of model training and inferring between normal pipeline and ColabSaprot. ColabSaprot streamlines the process, offering simplified procedures for model training and inference with just a few clicks. **b,** A lightweight plugin architecture (i.e., adapter) is integrated within Saprot to facilitate efficient training, sharing, and co-construction. Throughout this process, the model parameters of the Saprot backbone remain unchanged. **c,** ColabSaprot performs predictions by aggregating multiple shared models using adapters in SaprotHub without the need for private training data. **d,** Illustration of SaprotHub features, enabling biologists to co-share, co-build, co-use and collaborate within the community. **e,** ColabSaprot’s community-wide model collaboration mechanism (see **c**) allows it to achieve higher performance (orange bars) by aggregating multiple individually trained models (blue bars). Each individual model is trained with its own data, which may or may not overlap (reflecting real-world situations, see Supplementary Table 10). Data represents the mean ± s.d. of n = 5 independent experiments. **f,** By continually learning on models trained and shared by other biologists, ColabSaprot substantially outperforms those training-from-scratch models, especially when users lack sufficient training data (x-axis represents the number of training examples). Data represents the mean ± s.d. of n = 5 independent experiments. **g,** User study on supervised fine-tuning and zero-shot mutation effect prediction tasks (see Supplementary Table 11). For fine-tuning tasks, the data represents the mean ± s.d. of n = 5 independent experiments. **h,** User study on the inverse folding task. Experimental structures are shown in green, while predicted structures are in blue.

We first developed Saprot, a cutting-edge, large-scale protein language model that forms the foundation for ColabSaprot and SaprotHub. Saprot introduces a novel protein alphabet and representation, distinct from traditional amino acid (AA) sequences or explicit 3D coordinate structures. This alphabet is structure-aware (SA), with each “letter” encoding both the AA type and the local geometry of the protein. Formally, the SA alphabet (*SAA*) is defined as *SAA* = V × F, representing the Cartesian product of V and F, where V denotes the 20 AA and F represents the 20 structural (3Di) letters (Fig. 1a). These 3Di tokens are derived from protein 3D structures through Foldseek discretization [19]. The SA alphabet encompasses all possible combinations of AA (capital letters) and 3Di tokens (lowercase letters) in *SAA*, such as *Aa, Ap, Ac, …, Y a, Y p, Y y*, allowing proteins to be represented as a sequence of SA tokens that captures both primary and tertiary structures (Fig. 1b and Methods 1.1). Despite its conciseness, applying the SA alphabet successfully addresses the key challenges of scalability and overfitting in training on large-scale *AlphaFold2-generated* atomic structures (see Supplementary Note 1 and Supplementary Fig. 1). The adoption of “AA + structural token” sequences (Fig. 1b) for protein representation has garnered increasing attention in much subsequent studies [20, 21, 22, 23, 24, 25, 26, 27], emerging as a promising paradigm for protein representation.

The Saprot model uses a bidirectional Transformer [28] architecture (Fig.1b). It was pre-trained to reconstruct certain partially masked tokens in the SA token sequences (Methods 1.2). The model was trained from scratch on 40 million protein SA sequences, filtered from AlphaFold’s 214 million proteins [29] at 50% identity (Methods 1.3). Saprot is available in three sizes: 35M (million), 650M, and 1.3B (billion) parameters (Supplementary Table 2). The 650M model, used for most evaluations unless otherwise specified (see Supplementary Tables 3, 4), was trained for three months using 64 NVIDIA A100-80GB GPUs, representing computational resources comparable to those employed for ESM-2 (650M).

After its pre-training, Saprot has become a general-purpose foundation protein language model that excels across diverse protein prediction tasks, such as various supervised training tasks [30, 31, 32] (encompassing both regression and classification at either protein-level or residue-level), zero-shot mutation effect prediction [33, 34], and protein sequence design [35] (Fig. 1c, d and Supplementary Table 1, 5). Fig. 1d shows Saprot’s superior performance across 14 different protein prediction tasks compared to two well-known PLMs: ESM-2 [6] and ProtBert [9] (see detailed task, dataset, baseline and experimental setup descriptions in Methods 1.5, 1.8,1.9, 1.11, Supplementary Table 6, and more baseline comparison in Supplementary Table 7, 8). For tasks where protein structural information is available (indicated by purple, blue, and red colors), Saprot consistently surpasses both models. Even in scenarios without structural data (green tasks), Saprot maintains competitive performance with ESM-2 under the fine-tuning settings. Notably, Saprot substantially outperforms ESM-2 in three zero-shot mutational effect prediction tasks (Methods 1.6, 1.7): Mega-scale [36] (0.574 vs. 0.478), ProteinGym [37] (0.457 vs. 0.414), and ClinVar [38] (0.909 vs. 0.862) (see more results in Supplementary Table 7). Furthermore, despite being trained with a masked language modeling objective not specifically optimized for generation, Saprot performs effectively in protein sequence design while achieving a 16-fold acceleration in inference speed compared to ProteinMPNN [35](Fig.1e) (Methods 1.10, Supplementary Fig. 2 and Supplementary Table 9). Additionally, recent studies have demonstrated Saprot’s effectiveness across a broad spectrum of applications [39, 40], including protein engineering and de novo design [41, 42, 43], fitness and stability prediction [44, 45], molecular understanding [46, 47], fast & sensitive structure search [48], and drug-target interaction prediction [49, 50], etc. This versatility across multiple protein-related tasks supports SaprotHub’s vision for community collaboration (Supplementary Table 1).

We then developed the ColabSaprot platform by integrating Saprot into Google Colab’s infrastructure to support PLM training and prediction (Methods 1.14, Supplementary Fig. 3). ColabSaprot enables seamless deployment and execution of various task-specific trained Saprot models, eliminating the need for environment setup and code debugging. It also allows researchers to initiate training sessions with just a few clicks (Fig. 2a). ColabSaprot is designed to accommodate all tasks within the original Saprot framework, enabling direct prediction for tasks such as zero-shot mutation effect prediction [51, 34, 52] and protein sequence design [53, 35]. For mutation effect prediction, it implements single-site, multi-site, and whole protein sequence single-site saturation mutation. For protein design, it can generate de novo sequences based on a given backbone structure.

For these supervised training tasks, a particular focus of this work, users can fine-tune ColabSaprot with their own experimental data. Here, we implement a parameter-efficient fine-tuning technique [16, 17] by integrating lightweight adapter networks into ColabSaprot (Fig. 2b and Methods 1.12). During training, only adapter parameters are updated, achieving comparable accuracy as fine-tuning all Saprot parameters (Supplementary Fig. 4). This design not only improves learning efficiency [54, 55, 56, 57], but also establishes a collaborative and centralized framework that enables biologists to fine-tune task-specific Saprot within the research community, particularly through cloud-based environments (Fig. 2c,d,e,f). With adapters and the ColabSaprot interface, researchers can easily store and exchange their re-trained Saprot models in SaprotHub by loading or uploading adapter networks instead of the full pre-trained model. Since adapter networks contain much fewer parameters (around 1% of the whole Saprot model), this method greatly reduces storage, communication, and management burdens on SaprotHub, making model accessibility, co-construction, co-utilization, and co-sharing possible for a broader and diverse scientific community (see Fig. 2b,c,d and Methods 1.13).

Additionally, we developed several key features to streamline workflows for researchers. The ColabSaprot interface supports both sequence and structure inputs, offering automated dataset handling capabilities including efficient large-file upload mechanisms, real-time training monitoring with loss visualization, automatic saving and evaluation of the best checkpoints, break-point training resumption, and numerous safety checks to minimize user errors. To enhance GPU memory efficiency, we implemented adaptive batch sizing through gradient accumulation [4]. The platform also includes customizable settings, allowing researchers to modify code and adjust training parameters for specific research needs.

Finally, we developed SaprotHub with an integrated search engine, providing a centralized platform for sharing and collaboratively developing peer-retrained Saprot models within the biology community (Fig. 2d and Supplementary Fig. 5). Through the ColabSaprot interface, we implemented features like model storage, model sharing, model search, model continuous learning, and multiple model aggregation for enhanced performance (Methods 1.13,1.14). Specifically, SaprotHub offers three key advantages: First, researchers can share their trained models on SaprotHub for broader scientific applications without worrying about the leakage of private data. This feature effectively promotes knowledge dissemination and has the potential to establish a new paradigm for collaborative research (Fig. 2c). Second, researchers can leverage shared models on SaprotHub contributed by peers to perform continuous training on their own dataset [18]. This approach is particularly advantageous in data-limited scenarios, as fine-tuning from a better pre-trained model typically delivers superior predictive performance (Fig. 2f). Furthermore, these fine-tuned models can be contributed back to the hub through one-click upload of the adapter networks, fostering a collaborative ecosystem where the community collectively and iteratively enhances the available PLM resources. Third, as the SaprotHub community expands, it will accumulate a diverse collection of models for various protein prediction tasks, with multiple fine-tuned Saprot models becoming available for specific protein functions. To leverage this growing model collection, we implemented a model aggregation mechanism in ColabSaprot (see Methods 1.13, 1.14), enabling users to enhance predictive performance (Fig. 2e) through the integration of multiple existing models (Fig. 2c).

Saprot and SaprotHub have attracted community attention and demonstrated usefulness through multiple wet lab validations. The Zhang team at a commercial biological company employed ColabSaprot for the zero-shot single-point mutation prediction on a xylanase (NCBI ID: XP 069217686.1) from *Mycothermus thermophilus* and experimentally validated the top 20 predicted variants. Among these, 13 variants exhibited enhanced enzyme activity, with R59S showing a 2.55-fold improvement and F212N demonstrating a 1.88-fold increase in enzyme activity along with enhanced thermostability (see Methods 1.16, Supplementary Table 12, 13, 14 and Supplementary Fig. 6, 7). Similarly, the Chang lab employed ColabSaprot to perform zero-shot single-point mutation predictions on TDG, a uracil-N-glycosylase variant. Following experimental validation in HeLa cells, the top 20 variants were incorporated into the nCas9 protein. At the Dicer-1 target site, 17 out of the 20 predicted mutations showed enhanced editing efficiency compared to the wild type, as measured by the percentage of T-to-G substitutions at position T5 (the fifth thymine in the target sequence). Notably, three mutations - L74E, H11K, and L74Q - achieved nearly doubled editing efficiency (see Methods 1.18 and Supplementary Table 15). Another OPMC member recently fine-tuned Saprot using a dataset of approximately 140,000 GFP variants with corresponding fluorescence intensities to predict brighter avGFP variants from a pool of 5 million candidates. Experimental validation revealed that 7 out of the top 9 predicted double-site variants exhibited enhanced fluorescence compared to the wild-type, with one variant reaching more than 8-fold of the wild-type fluorescence intensity (see Methods 1.17 and Supplementary Table 16). Similarly, the Zheng lab shared an eYFP fluorescence prediction model, trained on 100,000 experimentally validated variants, achieved a Spearman correlation (*ρ*) of 0.94 with experimental fluorescence intensity on an independent test set of 6,000 variants, demonstrating near-experimental accuracy for double-site and triple-site mutations (see Model-EYFP 100K-650M on the SaprotHub webpage). We have recently received more feedback from community researchers who obtained positive wet lab results using ColabSaprot (see section 2).

We also conducted a user study by recruiting 12 biology researchers (without an ML background) and compared their performance with that of an AI expert (see details in Methods 1.15). The results demonstrated that, with ColabSaprot and SaprotHub, biology researchers can train and use state-of-the-art PLMs with performance comparable to that of an AI expert (Fig. 2g, h). Notably, in certain scenarios—such as the eYFP fitness prediction task illustrated in Fig. 2g—biologists leveraging pre-existing models from SaprotHub achieved higher prediction accuracy than AI experts. This higher performance stems from the fact that these shared models have been trained on larger or higher-quality datasets, highlighting the potential of model sharing within the scientific community—a key argument in this paper.

ColabSaprot and SaprotHub enable biology researchers to train and share sophisticated PLMs for diverse prediction tasks, even without extensive AI expertise. This platform empowers the broader protein research community to contribute and exchange PLMs, facilitating collaborative research and knowledge sharing through peer-trained models. We have made both Saprot and ColabSaprot open-source (see sections 2,3), providing a framework for other PLMs to develop their own model hubs. Importantly, ColabSaprot and SaprotHub represent just the first step in this evolution; our OPMC members have expanded this ecosystem by integrating more cutting-edge PLMs, including ProTrek [58] (35M, 650M), ESM-2 [6] (35M, 150M, 650M) and ESM-1b [5] (650M), and ProtBert (420M) [9] and ProtT5 (1.2B) [9] into the OPMC framework, thereby democratizing access to diverse PLMs for biologists worldwide (see section 3).

This community-wide participation approach to protein language modeling aligns with the OPMC vision. Our goal here is to inspire and foster the cooperative construction of open protein language models through SaprotHub. We envision SaprotHub as the catalyst that initializes OPMC, driving innovation and collaboration in the field.

## Acknowledgements

We thank Nan Li and Westlake University HPC Center for computing resources, and Jing Huang, Jianyang Zeng, Dangshen Li, Longxing Cao, Dapeng Li for discussions and paper advice. We thank John M. Jumper for providing insights about AF2 Evoformer and for his advice on the SA token design. This work is supported by the National Key Research and Development Program of China (No. 2022ZD0115100), the National Natural Science Foundation of China (No. U21A20427), the Westlake Center of Synthetic Biology and Integrated Bioengineering (WE-SynBio), Zhejiang Province Leading Geese Plan (No.2025C01094), the Zhejiang Key Laboratory of Low-Carbon Intelligent Synthetic Biology (No.2024ZY01025), and the Research Center for Industries of the Future (No.WU2022C030).

## Author contributions Statement

F.Y. conceived and led this research. J.S. conducted the main research and managed the technical implementation. J.S., F.Y., and J.S. (Junjie) designed the SA token and Saprot. J.S. and Z.L. developed the ColabSaprot and SaprotHub. T.T. developed the ColabSeprot and SeprotHub. F.Y., J.S., and S.O. designed the core idea of ColabSaprot and SaprotHub. S.O. and M.S. participated in discussions of ColabSaprot and SaprotHub, provided constructive ideas, and revised the manuscript. C.H. conducted partial zero-shot mutation experiments. F.D. trained the Saprot 1.3B version. Y.Z. collected the AFDB dataset. Y.H., X.C., & X.Z. (TDG), Q.Y. (xylanase), and Y.G. & T.S. (GFP) conducted the wet lab experiments. S.J. conducted some experimental result analysis. D.M. participated in early wet lab experiments. J.S. and F.Y. wrote the manuscript. Other OPMC authors contributed on wet lab validation, development of ColabSaprot and ColabSeprot, manuscript writing and proofreading, idea discussions, expert advice, promoting and testing ColabSaprot and ColabSeprot.

## Full author and affiliation lists

**The authors of this paper comprise all OPMC members listed below (in random order).** Anthony Gitter, University of Wisconsin-Madison, Madison, USA. Milot Mirdita, Seoul National University, Seoul, South Korea. Kevin K. Yang, Microsoft Research, Cambridge, USA. Pascal Notin, Harvard Medical School, Boston, USA. Debora S. Marks, Harvard Medical School, Boston, USA. Pranam Chatterjee, University of Pennsylvania, Philadelphia, USA. Rohit Singh, Duke University, Durham, USA. Philip A. Romero, Duke University, Durham, USA. Michael Heinzinger, Institute of Computational Biology, Neuherberg, Germany. Jianming Liu, Westlake University, Hangzhou, China. Jia Zheng, Westlake University, Hangzhou, China. Stan Z. Li, Westlake University, Hangzhou, China. Anping Zeng, Westlake University, Hangzhou, China. Huaizong Shen, Westlake University, Hangzhou, China. Jijie Chai, Westlake University, Hangzhou, China. Feng Ju, Westlake University, Hangzhou, China. Noelia Ferruz, Centre for Genomic Regulation (CRG), The Barcelona Institute of Science and Technology, Barcelona, Spain & Universitat Pompeu Fabra(UPF), Barcelona, Spain. Anum Glasgow, Columbia University, New York, USA. Philip M. Kim, University of Toronto, Toronto, Canada. Christopher Snow, Colorado State University, Fort Collins, USA. Vasilis Ntranos, University of California, California, USA. Jianyi Yang, Shandong University, Jinan, China. Liang Hong, Shanghai Jiao Tong University, Shanghai, China. Caixia Gao, Chinese Academy of Sciences, Beijing, China. Tong Si, Chinese Academy of Sciences, Shenzhen, China. Michael Bronstein, University of Oxford, Oxford, United Kingdom. Xing Chang, Westlake University, Hangzhou, China. Martin Steinegger, Seoul National University, Seoul, South Korea. Sergey Ovchinnikov, Massachusetts Institute of Technology, Cambridge, USA. Fajie Yuan, Westlake University, Hangzhou, China. Jin Su, Westlake University, Hangzhou, China. Zhikai Li, Westlake University, Hangzhou, China. Tianli Tao, Westlake University, Hangzhou, China. Chenchen Han, Westlake University, Hangzhou, China. Yan He, Westlake University, Hangzhou, China. Fengyuan Dai, Westlake University, Hangzhou, China. Xuting Zhang, Westlake University, Hangzhou, China. Yuyang Zhou, Westlake University, Hangzhou, China. Junjie Shan, Westlake University, Hangzhou, China. Xibin Zhou, Westlake University, Hangzhou, China. Shiyu Jiang, Westlake University, Hangzhou, China. Dacheng Ma, Zhejiang Lab, Hangzhou, China. Yuan Gao, Chinese Academy of Sciences, Shenzhen, China. Jiawei Zhang, Westlake University, Hangzhou, China. Yuliang Fan, Westlake University, Hangzhou, China. Yuyang Tao, ShanghaiTech University, Shanghai, China. Linqi Cheng, Rice University, Houston, USA. Xinzhe Zheng, Rice University, Houston, USA. Lei Chen, Shenzhen Lions King Hi-Tech Co., Ltd, Shenzhen, China. Rui Long, Shenzhen Lions King Hi-Tech Co., Ltd, Shenzhen, China. Lingjie Kong, South China Agricultural University, Guangzhou, China. Zhongji Pu, Xianghu laboratory, Hangzhou, China. Jiaming Guan, Hefei MiQro Era Digital Technology Co. Ltd., Hefei, China. Tianyuan Zhang, Suzhou Polynovo Biotech Co., Ltd., Suzhou, China. Cheng Li, Suzhou Polynovo Biotech Co., Ltd., Suzhou, China. Qingyan Yuan, Suzhou Polynovo Biotech Co., Ltd., Suzhou, China.

## 1 Methods

### 1.1 Constructing SA protein sequence

The Structure-Aware (SA) vocabulary encompasses both residue and structure information, as illustrated in Fig. 1**a**. Given a protein *P*, its primary sequence can be denoted as (*s*_1_*, s*_2_*, …, s_n_*), where *s_i_* ∈ V represents the residue at the *i_th_* site, and V represents residue alphabet. Drawing inspiration from the vector quantization learning technique [81], we encode protein 3D structures into discrete residue-like structural tokens. Here, we use Foldseek [19], a fast and accurate protein structure aligner. Through Foldseek, we have a structure alphabet F, wherein *P* is expressed as the sequence (*f*_1_*, f*_2_*, …, f_n_*), with *f_j_* ∈ F representing the 3Di token for the *j_th_* residue site. To maintain simplicity, we adhere to the default configuration of Foldseek, which sets the size *m* of F to 20. We then combine the residue and structure tokens per residue site, generating a new SA protein sequence *P* = (*s*_1_*f*_1_*, s*_2_*f*_2_*, …, s_n_f_n_*), where *s_i_f_i_* ∈ V × F is the so-called SA token naturally fusing both residue and geometric conformation information. The SA-token protein sequence can then be fed into a standard Transformer encoder as basic input. It’s important to note that we also introduce a mask signal “#” to both residue and structure alphabet, which results in “*s_i_*#” and “#*f_i_*” that indicate only residue or structure information is available. The size of the SA vocabulary is 21 × 21 = 441.

### 1.2 Model architecture and pre-training of Saprot

Saprot follows the same network architecture and parameter size as ESM-2 [6], which draws inspiration from the BERT [28] model in Natural Language Processing (NLP). The primary difference lies in the embedding layer: Saprot incorporates 441 SA tokens in place of the conventional 20 amino acid tokens. This nearly identical architecture enables straightforward performance comparison with the ESM model.

Saprot is pre-trained with the Masked Language Modeling (MLM) objective [28], like ESM-2 and BERT. Formally, for a protein sequence *P* = (*s*_1_*f*_1_*, s*_2_*f*_2_*, …, s_n_f_n_*), the input and output can be represented as: *input*: (*s*_1_*f*_1_*, …,* #*f_i_, …, s_n_f_n_*) → *output*: *s_i_f_i_* (Fig. 1b). Given that the 3Di token may not always be accurate for certain regions in predicted structures by AlphaFold2, *f_i_* in #*f_i_* is made visible during training so as to reduce the emphasis the model places on its predictions.

AlphaFold2 (AF2) predictions include pLDDT confidence scores that indicate the precision of predicted atom coordinates. Therefore, we implement specialized handling for regions with low confidence scores. During pre-training, regions with pLDDT scores below 70 are processed distinctly. When these regions are selected for MLM prediction, we use the “*s_i_*#” token as the prediction target, while masking the input SA sequence with the “##” token. This approach encourages the model to predict residue types based solely on contextual information. When these low-confidence regions are not selected for MLM prediction, we employ the “*s_i_*#” token in the input, ensuring the model relies solely on residue context rather than inaccurate structural information. For downstream tasks, we maintain consistency with the pre-training protocol by applying the same handling to regions with pLDDT scores below 70. These regions are represented using “*s_i_*#” tokens, with only residue information remaining visible.

Saprot 35M and 650M underwent typical pre-training from scratch as described above. In contrast, Saprot B employed an efficient training strategy [69] by architecturally combining two identical 33-layer Saprot 650M models. The initialization process involved duplicating the pre-trained 650M model’s parameters to populate both the lower (layers 1-33) and upper (layers 34-66) sections of the 1.3B architecture. After initialization, Saprot 1.3B was trained following the same training protocol employed for the 35M and 650M models, except that 30% of the protein sequences in a training batch were transformed into the amino acid sequence only format (*s*_1_#*, s*_2_#*, …, s_n_*#) to enhance the model’s ability to handle proteins without available structural information. Training was terminated upon convergence of the loss function.

### 1.3 Processing pre-training dataset

We followed the procedures outlined in ESM-2 [6] to generate sequence identity filtered protein data. Subsequently, we acquired all AF2 structures via the AlphaFoldDB website (https://alphafold.ebi.ac.uk/), utilizing the UniProt IDs of protein sequences. Proteins without structures in AlphaFoldDB were removed. This process yielded a collection of approximately 40 million structures. Employing Foldseek, we encoded these structures into 3Di tokens. Subsequently, we formulated SA sequences by combining residue and 3Di tokens, treating them as a single SA token at each position. These datasets were used for training all three versions of the Saprot models.

### 1.4 Hyper-parameters for pre-training

Following ESM-2 and BERT, during training, 15% of the SA tokens in each batch were masked. We replaced the SA token *s_i_f_i_* with the #*f_i_* token 80% of the time, while 10% of the tokens were replaced with randomly selected tokens, and the other 10% tokens remained unchanged. For the optimization of Saprot, we adopted similar hyper-parameters to those employed in the ESM-2 training phase. Specifically, we employed the AdamW optimizer [59], setting *β*_1_ = 0.9*, β*_2_ = 0.98 and we utilized *L*_2_ weight decay of 0.01. We gradually increased the learning rate from 0 to 4e-4 over the first 2000 steps and linearly lowered it to 4e-5 from 150K steps to 1.5M steps. The overall training phase lasted approximately 3M steps. Like the ESM model, we also truncated them to a maximum of 1024 tokens, and our batch size consisted of 512 sequences. Additionally, we also employed mixed precision training to train Saprot.

### 1.5 Descriptions of baseline models

We compared Saprot to several prominent protein language models (see Supplementary Table 7, 8). For supervised learning tasks, we compared Saprot with ESM-2 (the 650M version) [6], ProtBert (the BFD version) [9], MIF-ST [82], GearNet [60] and ESM-3 [20]. The first two models utilize residue sequences as input, while the latter three models incorporate both residue and structures as input. ESM-2 (650M) stands out as the primary baseline for comparison, given its similar model architecture, size, and training approach when compared to Saprot. ESM-2 also offers a 15 billion (B) parameter version, which can be challenging to fine-tune even on GPUs with 80G memory. Therefore, we only conducted comparisons with ESM-2 (15B) for zero-shot mutational effect prediction tasks, which can be achieved without the need for fine-tuning.

For the zero-shot mutational effect prediction task, we compare with the state-of-the-art ESM-2, ProtBert, ESM-1v [7], Tranception L (without MSA retrieval) [11], MSA Transformer [8], EVE [34] and ESM-3 models. For the protein inverse folding task, we compare with ProteinMPNN [35] as baseline.

### 1.6 The formula for the zero-shot mutation effect prediction task

Previous sequence-based protein language models like the ESM models predict mutational effects using the log odds ratio at the mutated position. The calculation can be formalized as follows:

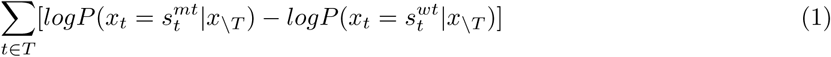

Here *T* represents all mutations and *s_t_* ∈ V is the residue type for mutant and wild-type sequence. We slightly modify the formula above to adapt to the SA alphabet in Saprot, where the probability assigned to each residue corresponds to the summation of tokens encompassing that specific residue type, as shown below:

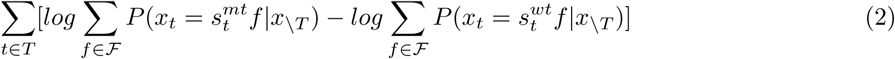

Here *f* ∈ F is the 3Di token generated by Foldseek and *s_t_f* ∈ V × F is the SA token in our new alphabet.

### 1.7 Zero-shot mutational effect prediction tasks and datasets

#### ProteinGym

ProteinGym [37] comprises an extensive collection of Deep Mutational Scanning (DMS) assays, enabling comprehensive comparison among zero-shot predictors. We evaluate all baseline models (see Supplementary Table 7) on ProteinGym’s substitution branch, utilizing its provided protein structures and adhering to the standard evaluation protocol outlined in the original paper [37]. We employ Spearman’s rank correlation as our evaluation metric.

#### ClinVar

ClinVar [38] is a publicly accessible repository housing information about human genetic variants and their clinical significance in disease. For our analysis, we utilize data curated from EVE [34], excluding proteins exceeding 1024 residues in length. To ensure data quality, we restrict our analysis to proteins with reliability ratings of 1 “Gold Star” or higher. Following EVE’s methodology, we assess model performance using the AUC metric.

#### Mega-scale

Mega-scale [36] employs cDNA display proteolysis to measure thermodynamic folding stability across protein domains. The dataset encompasses all single mutations and selected double mutants in both natural and *de novo* designed proteins. For this dataset, we also utilize Spearman’s rank correlation as our evaluation metric.

For each mutation dataset, we provide all variants with the wild-type structure, as AF2 does not reliably distinguish the structural changes induced by single mutations. Additionally, the ClinVar dataset only provides UniProt IDs, so we manually downloaded all AF2 structures and eliminated proteins without structures in AlphaFold DB. Both ProteinGym and Mega-scale datasets provide protein structures, either predicted from AF2 or derived from *de novo* design.

### 1.8 Supervised fine-tuning tasks and datasets

#### Fine-tuning Saprot with the AF2 structure

These benchmarks include the Thermostability task from FLIP [61] and the Localization Prediction task from DeepLoc [62]. The DeepLoc benchmark has two prediction branches: a multi-class classification with 10 subcellular locations and a binary classification with 2 location categories. We use the datasets provided by these original literature. For structural information, we obtain AlphaFold2-predicted structures for all proteins using their corresponding UniProt IDs.

#### Fine-tuning Saprot with the PDB structure

There are a few tasks providing experimentally determined structures as training data. We evaluate Metal Ion Binding task [72] and a series of tasks from ProteinShake [31], including Structure Class Prediction, Structural Similarity Prediction and Binding Site Detection. The corresponding datasets are provided in the respective literature.

#### Fine-tuning Saprot without structure

While Saprot is designed to leverage protein structural information, it can still work in scenarios where structural data is not provided during supervised fine-tuning (also see Methods 1.2). In these cases, we mask the 3Di token in the SA sequence. We evaluate performance on the Fluorescence and Stability prediction datasets from TAPE [30], the AAV dataset from FLIP [61], and the *β*-lactamase landscape prediction dataset from PEER [32].

Regarding the evaluation metrics, we adopt those established in the original literature. For details on dataset splits, please refer to the next subsection.

### 1.9 Data split

In existing literature, datasets are typically partitioned based on protein sequence identity. However, a recent benchmark study called ProteinShake [31] argued that protein structures exhibit higher conservation than sequences, indicating that structure-based splitting provides a more stringent evaluation of model generalization. Inspired by this, we have adopted the same structure-based data splitting for most of our evaluation (unless otherwise specified). The splitting criterion is quantified using the Local Distance Difference Test (LDDT), as proposed in ProteinShake. To be specific, for datasets that include protein structures (Methods 1.8), we employ the default 70% LDDT threshold recommended in ProteinShake [31]. For tasks where only sequence data is available, we retain the original splits provided in the official literature, as these datasets consist of only mutational variants of one single protein. To show performance sensitivity across different splitting criteria, we perform further evaluation by comparing Saprot (35M) and ESM (35M) using a more stringent 30% LDDT threshold and the commonly used 30% sequence identity threshold [32, 63]. The corresponding results are presented in Supplementary Table 3.

### 1.10 Protein inverse folding

#### Saprot for protein inverse folding

To use Saprot for inverse folding, we first encode the protein backbone into 3Di tokens (*f*_1_*, f*_2_*, …, f_n_*). We then mask all residue parts of the SA tokens, forming an SA sequence (#*f*_1_, #*f*_2_*, …,* #*f_n_*). This sequence is input into Saprot to predict residue distributions at all positions. In contrast to ProteinMPNN [35] that generates residues in the auto-regressive manner, i.e. generating next token conditioned on all previous outputs, Saprot is able to simultaneously predict all residues with only one forward propagation.

#### Fine-tuning Saprot on the CATH Dataset

We evaluate two variants of Saprot for protein design: one pre-trained using AF2-predicted structures and another further fine-tuned on the CATH dataset [76]. This CATH database, which was also used to train ProteinMPNN [35], is partitioned into training, validation, and test sets using an 80/10/10 split, following Ingraham et al. [73]. For detailed dataset construction procedures, readers are referred to [73].

### 1.11 Hyper-parameters for fine-tuning tasks

We employ the AdamW [59] optimizer during fine-tuning, setting *β*_1_ = 0.9*, β*_2_ = 0.98, along with *L*_2_ weight decay of 0.01. We use a batch size of 64 for all datasets. We empirically found that the optimal learning rate for most baselines are in the range of 1e-5 to 5e-5. For **Training with AF2 structure** and **Training with PDB structure**, the optimal learning rate was set to 5e-5, whereas for **Training without structure**, it was set to 1e-5. We fine-tuned all model parameters until convergence and selected the best checkpoints based on their performance on the validation set.

### 1.12 Adapter learning

The adapter learning technique, originated in general AI community [71, 67, 83], has recently been adopted in protein research [56, 54], [78, 77]. While these studies mainly focused on its advantage for model accuracy and training efficiency, here we use adapter by integrating it with Google Colab to create a platform that enables model fine-tuning, sharing, continuous re-training, and collaboration within the biological research community through our online SaprotHub.

In the broader AI community, various types of adapters exist, including Houlsby [71], Pfeiffer [17], Compacter [74], and LoRA [16]. We choose LoRA for its capacity to deliver comparable results with fewer parameters. By integrating learnable low-rank matrices into each Saprot block while freezing the backbone, LoRA enables parameter-efficient fine-tuning and model sharing for these downstream tasks.

### 1.13 Community-wide collaboration

As the SaprotHub community grows, more fine-tuned Saprot models become available for each protein function, facilitating collaborative development among biologists. To evaluate this advantage (see Fig. 2e), we adopt the following approach: For each task, we randomly partition the training data into five subsets and train one model on each subset, yielding models model 0 through model 4, simulating models shared by different researchers. For regression tasks (e.g., Thermostability), the final prediction (model agg) is computed as the mean of all model outputs. For classification tasks (e.g., Subcellular Localization), we implement majority voting, where the final prediction (model agg) is determined by the most frequent class prediction across the ensemble. The lightweight adapter technique is crucial for model aggregation, addressing the challenges of loading and sharing multiple large pre-trained models.

To evaluate the effectiveness of model continue learning (Fig. 2f), we randomly sampled 100, 200, and 500 training instances per task to simulate researchers’ private data. We then performed fine-tuning using two base models, updating only their adapter parameters: the official Saprot model (blue) and a shared model from SaprotHub (orange). The latter represents models previously trained by researchers with larger or higher-quality datasets. Results demonstrate that continuing training from existing well-trained models substantially outperforms training from scratch with limited data. Note that the official Saprot model (blue) is pretrained on large-scale protein sequence and structure data but has not undergone supervised fine-tuning for specific tasks. This highlights a key advantage of SaprotHub: researchers can build upon others’ achievements.

### 1.14 ColabSaprot notebooks

ColabSaprot consists of three key components, with the first focusing on model training. This component enables researchers to rapidly configure the runtime environment, process training data, and fine-tune Saprot efficiently. We provide access to four base models: Saprot-35M, Saprot-650M, custom fine-tuned Saprot models, community-shared fine-tuned models from SaprotHub. Additionally, we offer advanced hyper-parameter customization options, allowing researchers to tailor their training strategies to specific requirements. These customizable parameters include batch size, learning rate, and training steps etc.

The second component focuses on prediction capabilities. ColabSaprot supports a diverse range of prediction tasks (see Supplementary Table 1) using both community-shared models from SaprotHub and locally fine-tuned models. This includes protein-level, residue-level, and protein-protein interaction predictions, as well as zero-shot mutational effect predictions and protein design. Each task category offers multiple configuration options to accommodate different research requirements. Moreover, ColabSaprot enhances prediction accuracy by implementing ensemble methods that aggregate multiple models from SaprotHub, delivering more robust and reliable results. This collaborative approach facilitates a community-driven model ecosystem where researchers can leverage and combine multiple models to achieve superior performance.

The third component enables model sharing, searching (implemented in SaprotHub, see Supplementary Fig. 5) and community collaboration. Upon completion of training, researchers can contribute their model weights (specifically the adapter weights) to the SaprotHub community repository, making them accessible to the broader biological research community. Through SaprotHub’s specialized search engine, researchers can efficiently locate and utilize relevant models. These shared models can be leveraged for continual learning, direct application, or model aggregation to achieve enhanced performance.

### 1.15 User study

We evaluated ColabSaprot-v1 through eight protein prediction tasks with twelve participants who, based on brief conversations and background checks (see detailed material through the provided link in Data Availability section), had biological backgrounds but no prior involvement in coding or machine learning projects. The evaluation encompassed supervised fine-tuning, zero-shot mutation effect prediction, and protein inverse folding tasks. Each participant had three days to complete their assignments (task descriptions are attached through the provided link in Data Availability section) and received compensation upon submitting either their results or documentation of encountered obstacles via screenshots or recordings.

For comparison, we engaged an AI expert - a third-year PhD student specializing in machine learning with over two years of research experience in protein-focused AI applications. The expert executed these tasks using the Saprot codebase from our GitHub repository, conducting five independent runs with optimized hyperparameters using different random seeds.

The twelve participants were organized as follows: one was assigned to zero-shot mutational effect prediction, another to protein design, and the remaining ten were randomly divided into two groups of five. Each group handled three supervised fine-tuning tasks, covering six tasks in total. In other words, each supervised fine-tuning task was completed by 5 participants. We evaluated the average accuracy across all participants, including those who were unable to finish the task completely (when such cases occurred).

For the supervised fine-tuning tasks, we used five public datasets for predicting: thermostability, subcellular and binary localization, GFP fluorescence, and stability (see Supplementary Table 6). We also included a proprietary eYFP fitness prediction dataset from Zheng’s lab, containing 3,087 validation and 3,088 test samples, bringing the total to six supervised fine-tuning tasks. To reduce training time and computing power consumption, we randomly selected 1,000 samples from the training set of each dataset while keeping the validation and test sets unchanged.

For model evaluation, all participants generated predictions on designated test sets. In the eYFP task, biology participants leveraged continual learning on a pre-trained model from SaprotHub (Model-EYFP-650M (search model name via https://huggingface.co/spaces/SaProtHub/SaprotHub-search), achieving 0.95 Spearman’s *ρ* on test data), while the AI expert used the base Saprot 650M model. This setup is used to show how SaprotHub enables researchers with limited data to build upon existing fine-tuned models. For other tasks, both biology participants and the AI expert trained their models using the same SaProt 650M AF2 base model with identical training and test sets, ensuring fair comparison.

For the zero-shot mutational effect prediction, we randomly selected four mutation datasets from ProteinGym benchmark [37] for evaluation. Three of these datasets focus on the impact of mutations on enzyme activity, while the fourth addresses drug resistance. We assigned one participant to conduct predictions on these datasets using ColabSaprot in a zero-shot manner.

For the protein inverse folding task, we assigned one participant to use ColabSaprot to generate protein sequences based on given structures. Subsequently, the participant employed ESMFold (an interface provided by ColabSaprot) to predict the structures of the generated sequences. To assess Saprot’s ability on new proteins, we selected these recently released structures (see Fig. 2h).

More materials about participants, task descriptions and result evidence are available via link in the Data Availability section. Participants were provided with related protein datasets (including training and test sets), GPU-enabled Google Colab accounts, and detailed task instructions. Participants had three days to complete the assignments on their own using ColabSaprot. To encourage honest feedback and thorough documentation, compensation was guaranteed to all participants who documented their challenges through screenshots or recordings, regardless of whether they completed the assigned tasks.

We acknowledge a potential self-selection bias, as the biology participants were volunteers likely interested in novel computational tools. This may imply that their aptitude for learning new software could be higher than the general average for the biology community. However, as the study’s primary goal was a comparative analysis of workflows, this bias is not expected to alter the main conclusions regarding the platform’s relative efficiency and accessibility. All participants were fully informed about the study.

### 1.16 Experimental validation on the xylanase

There are several steps. Let’s name the protein (NCBI ID: XP 069217686.1) Mth first:

(1) Use AlphaFold3 (https://golgi.sandbox.google.com/) to get the protein structure of Mth;
(2) Use Foldseek or ColabSaprot to get the SA sequence of Mth;
(3) Use ColabSaprot (650M) to perform zero-shot (single-point) mutation effect prediction for the entire SA sequence;
(4) Choose the 20 highest-ranked variants (see Supplementary Table 12) excluding A15G and I13P; A15G and I13P were not selected because they are located in the signal peptide region of the protein, which would be removed during the process of protein secretion in *Pichia pastoris*.

(5) Construct mutants. See below for details:

(5.1) The gene sequence of Mth was optimized according to the codon preference of *P. pastoris*, and the plasmid pPIC9K-Mth was synthesised by GenScript (Nanjing, China).

(5.2) Mutations of Mth were generated by site-directed mutagenesis via PCR, using the plasmid pPIC9K-Mth as a template. The primers are listed in Supplementary Table 13.

(5.3) After confirmation by DNA sequencing, the wild-type and mutated plasmids were linearized with *Sal* I and used to transform *P. pastoris* strain GS115. The recombinant strains were selected on MD plates (1.34% YNB (yeast nitrogen base without amino acids), 2% glucose, 2% agar) and verified by PCR and sequencing.

(6) Enzyme activity assay of wild-type Mth and mutants. See below for details:

(6.1) The positive transformants were cultivated in liquid medium BMGY [1% yeast extract, 2% peptone, 1.34% YNB, 1% glycerol, 100 mM potassium phosphate (pH 6.0)] for 20-24 h. The cells were collected by centrifugation at 3500 x g and 4°C for 5 min, then transferred to 250 mL shake flasks containing 25 mL BMMY [1% yeast extract, 2% peptone, 1.34% YNB, 1% methanol, 100 mM potassium phosphate (pH 6.0)] with initial OD_600_ =0.5. Fed-batch fermentation was proceeded to express xylanase by feeding 1% methanol per 24 h. All liquid cultures were performed at 30°C and 250 rpm. After 120 h of cultivation, the supernatants were harvested by centrifugation at 3500 x g and 4°C for 5 min and tested for xylanase activity. (6.2) Xylanase activity assay: The reaction mixture contained 0.1 mL of 1% (w/v) beechwood xylan and 0.1 mL of a suitably diluted enzyme solution (100 mM acetate buffer, pH 5.0) incubated at 60°C for 30 min. The amount of reducing sugar released was determined using the 3,5-dinitrosalicylic acid (DNS) method, with xylose as the standard. One unit (U) of xylanase activity was defined as the amount of enzyme that catalyzes the release of 1 µmol of xylose equivalent per minute under the assay conditions. The enzyme activity was measured at 40–70°C to determine the optimal temperature of the enzymes, with the pH of the reaction maintained at 5.0. After incubation at 60°C for various time periods, the residual enzyme activity was measured to assess the thermostability of the enzymes.

### 1.17 Experimental validation on GFP variants

The objective of this task was to engineer brighter avGFP variants as part of the 2024 Critical Assessment of Protein Engineering (CAPE) competition [68]. CAPE, a student-focused challenge modeled after the Critical Assessment of Structure Prediction (CASP) competition, emphasizes protein function design and variant effect prediction. The parent GFP sequence is based on avGFP (see the amino acid sequence information in Supplementary Table 14) derived from Aequorea victoria (UniProtKB: P42212). These top-ranked predicted variants by Saprot were experimentally validated by the CAPE organizers.

The process of fine-tuning Saprot (35M) and subsequent prediction consisted of the following steps: Step 1: Data Preparation: The CAPE organizers provided a dataset of 140,000 GFP variants (including avGFP, amacGFP, cgreGFP, and ppluGFP2) (see Supplementary Table 14) along with their corresponding fitness scores, and the structures of four wild type GFP proteins (avGFP, amacGFP, cgreGFP, and ppluGFP2). Our OPMC member Xuting Zhang utilized Foldseek to generate 3Di tokens for these wild type structures, which were then used to converted to SA tokens by their corresponding variants.

Step 2: Model Fine-tuning: The Saprot model underwent full parameter fine-tuning using the SA token sequences of these variants.

Step 3: Variant Prediction and Validation: A pool of 5 million avGFP double-site mutants was generated through random mutagenesis. The fine-tuned Saprot model from Step 2 was used to predict their fitness scores. The top-9 ranked variants were selected for experimental validation.

Experimental validation was conducted using the robotic biofoundries at the Shenzhen Infrastructure for Synthetic Biology, following the below procedure:

Step 1: Expression plasmid pET28a vectors containing designed mutant GFP sequences were ordered from Genescript Biotech (Nanjing, China) and used to transform chemically competent E. coli BL21(DE3) cells through heat shock at 42°C. Two independent clones for each mutant were randomly selected to inoculate 1 mL of non-inducing LB+Kan medium (supplemented with 50 µg mL-1 kanamycin for plasmid maintenance by antibiotic selection) in 96-well microtiter plates to prepare seed cultures, which were grown at 37°C for 16-20 h.

Step 2: For inducible protein expression, 40 µL of stationary-phase seed cultures were used to re-inoculate 4 mL of fresh LB+Kan medium in 24-well microtiter plates, followed by approximately 4-h incubation at 37°C to reach the exponential growth phase (OD_600_=0.6-0.8). Then, IPTG was added to achieve a final concentration of 1 mM for inducible expression at 18°C for 20 h.

Step 3: Biomass growth was monitored by using OD_600_ measurement and GFP fluorescence was assessed with excitation at 488 nm and emission at 520 nm in 96-well flat-bottomed plates. For each strain, GFP signal intensities was divided by OD_600_ values to calculate biomass-normalized fluorescence.

### 1.18 Experimental validation of the TDG variants

The Chang lab utilized ColabSaprot v1 (650M) for zero-shot prediction of single-site mutation effects of a uracil-N-glycosylase variant (TDG) (see Supplementary Table 14 for its amino acid sequence). They input only the amino acid sequence with all structural tokens masked into ColabSaprot to identify the top 20 highest-ranked variants. Experimental validation and results of these variants were documented in Supplementary Table 15.

Following protocols described in [64, 51], base editors used in this paper were cloned in a pCMV plasmid with blasticidin resistance. sgRNA was cloned into a pSuper-sgRNA plasmid with puromycin resistance. The TDG sequence was amplified from plasmid TSBE2 [51], and TDG variants were generated via site-directed mutagenesis by PCR, then fused with SpCas9 (D10A) protein as TSBE2. All primers were listed in Supplementary Table 13. Protein sequence of TSBE2 and protospacer sequences of sgRNA are available in reference [51]. The experimental validation process was performed as previously described [64, 51].

## 2 Data availability

The pre-training dataset for training Saprot is available at https://huggingface.co/datasets/westlake-repl/ AF2 UniRef50. Downstream task datasets are all stored at https://huggingface.co/SaProtHub. Materials for user study are available at https://drive.google.com/file/d/1LdGRnwt2lttnszNBAJ0F967A8rguPq8b/view?usp=sharing. The raw data with detailed wet lab information is available at https://drive.google.com/file/d/1IYcOqRuF76LusG7DMI4kEr2mJL6aE7g-/view?usp=sharing. Additional wet lab experimental results collected from the research community are available at https://drive.google.com/file/d/1ZcDl0XYksTcUEaCfPI0tuEVo5CA031dg/view?usp=sharing. All unique/stable reagents generated in this study are available from the lead contact.

## 3 Code availability

Saprot is an open-sourced model with MIT license. The code is available at https://github.com/westlake-repl/Saprot. The code implementation of ColabSaprot notebook is available at https://github.com/westlake-repl/SaProtHub. ColabSaprot service (latest version: v2) is available at https://colab.research.google.com/github/westlake-repl/SaprotHub/blob/main/colab/SaprotHubv2.ipynb?hl=en (the previous v1 version is still maintained in the our SaprotHub github webpape). All fine-tuned Saprot models can be obtained through SaprotHub https://huggingface.co/SaProtHub by the dedicated search engine https://huggingface.co/spaces/SaProtHub/SaprotHub-search.

Our OPMC members have also implemented ColabSeprot (https://colab.research.google.com/github/westlake-repl/SaprotHub/blob/main/colab/ColabSeprot.ipynb?hl=en), including ColabProTrek (35M,650M), ColabESM1b (650M), ColabESM2(35M,150M,650M) and ColabProtBert(420M) and SeprotHub (https://huggingface.co/SeprotHub) and an independent ColabProtT5 (https://colab.research.google.com/github/westlake-repl/SaprotHub/blob/main/colab/ColabProtT5.ipynb).

## Supplementary Information OPMC

The Open Protein Modeling Consortium (OPMC) is a collaborative initiative aimed at bringing together the efforts of the protein research community. Its mission is to foster the sharing and co-development of resources, with an emphasis on individually trained decentralized models, helping to advance protein modeling through collective contributions. OPMC provides platforms/tools that support diverse protein prediction tasks, striving to make advanced protein modeling more accessible to researchers, regardless of their level of expertise in machine learning.

Here are some Q&A related to OPMC that we have gathered from our steering committee members and other researchers: https://theopmc.github.io/

### Supplementary Note 1

To realize the vision of community-wide collaboration at SaprotHub, the protein language model (PLM) Saprot should have both advanced performance and support for a diverse range of tasks. This necessitates that its pre-trained representations achieve a high degree of generalizability. Contrastive learning (CL) and BERT-style masked language modeling (MLM) are currently the two predominant pre-training approaches for generalized representation learning, however, CL primarily focuses on protein-level representation learning with limited effectiveness in residue-level tasks. Therefore, we decided to adopt the BERT-style MLM loss for training Saprot. However, a problem arises when directly training the AF2-predicted structure using the MLM loss. Here, we demonstrate this issue by applying two common methods (see Supplementary Fig. 1) to train PLMs on AF2-predicted structures with the MLM loss function.

Method 1: following MIF [82], we represented AF2-predicted structures as graphs and utilized Graph Neural Networks (GNNs) for modeling. Note that the original MIF model was pre-trained using only experimental structures, so the effectiveness of incorporating a large number of AF2 predicted structures remains uncertain.

Method 2: we extracted pairwise residue distance and angle information from structures to incorporate as structural bias in the Transformer attention maps, an approach previously implemented in Uni-Mol [84] and Evoformer. Also note that the two models were originally designed for different tasks with different loss functions, and here we would like to evaluate their performance using the AF2-predicted structures and the MLM loss.

The training loss curves are shown in Supplementary Fig. 1. Our experimental results demonstrate that PLMs directly trained on AF2-predicted structures by using the MLM loss (i.e., the two models on the left side in Supplementary Fig. 1) yields poor performance on validation sets containing experimental structures. Notably, while the loss decreases on predicted structures during training, we observe no corresponding improvement in performance on experimental structures. This discrepancy suggests that the PLM may have detected some potential traces or bias from AF2 predictions rather than genuine structural features. Consequently, despite the apparent decrease in training loss, the model fails to acquire meaningful representations of real protein structures. By contrast, the Saprot model show consistent loss decrease in both AF2-predicted structures and experimental structures. More details were reported in the ICLR version of our Saprot paper, see [14] Supplementary Table 8 and section 3.1.

These results explain why we considered training Saprot with discrete structural token rather than explicit 3D structures. A similar idea was recently adopted by ESM-3 [20], further demonstrating that the use of discrete structural token is an effective way to scale PLMs with a large number of structures. By contrast, existing PLMs [66, 70] that incorporate explicit 3D structural information often utilize only a small number of protein structures during pre-training, and have not demonstrated that scaling with more AF2-predicted structures improves performance, in particular with the MLM loss function.

---

**Supplementary Table 1:** Tasks that Saprot potentially support. As a large-scale foundation protein language model, Saprot potentially supports hundreds of protein prediction tasks (also see Fig. 1 c,d). These tasks include, but are not limited to: protein-level classification, protein-level regression, residue-level classification, protein-protein classification, protein-protein regression, zero-shot mutational effect prediction, and protein sequence design.

| Supported Tasks |  |
| --- | --- |
| **Protein-level Classification** | Active Site Prediction<br>Post-Translational Modification (PTM) Site Prediction<br>Mutation Impact Prediction<br>Transmembrane Region Prediction<br>Conservation Score Prediction<br>Interface Residue Prediction<br>Enzyme Binding Site Prediction<br>Metal Binding Site Prediction<br>Disease-Associated Variant Prediction<br>Immune Receptor Binding Prediction<br>Glycosylation Site Prediction |
| Fold Class Prediction<br>Localization Prediction<br>Subcellular Localization<br>Domain Classification<br>Membrane Protein Classification<br>Enzyme Classification<br>Post-translational Modification Site Classification<br>Protein Stability Classification<br>Protein Solubility Classification<br>Antigenicity Classification<br>Protein Aggregation Classification<br>Protein Family Classification<br>Protein Structural Class Prediction<br>Protein Subunit Interface Classification<br>Protein Motif Classification<br>Protein Epitope Classification<br>Protein Interaction Network Classification<br>Protein Toxicity Classification<br>Protein Expression Level Classification<br>Protein Dynamics Classification<br>Protein Flexibility Classification<br>Protein Function Annotation<br>Protein Interaction Type Classification<br>Protein Disease Association Classification<br>Protein Functional Domain Classification | **Protein-protein Classification** |
|  | Protein-Protein Interaction (PPI) Prediction<br>Interaction Type Classification<br>Disease-Associated Interaction Prediction<br>Mutational Impact Classification<br>Functional Annotation Classification<br>Interaction Network Module Detection<br>Interaction Stability Prediction |
|  | **Protein-protein Regression** |
|  | Interaction Strength Prediction<br>Binding Free Energy Calculation<br>Interaction Affinity Prediction<br>Mutational Impact Quantification<br>Interaction Kinetics Prediction |
|  | **Zero-shot mutational Effect prediction** |
| **Protein-level Regression** | Enzyme Activity Prediction<br>Virus Fitness Prediction<br>Driver Mutation Prediction<br>Protein Stability Prediction<br>Immune Escape Prediction<br>MHC Binding Prediction<br>Pathogenicity Prediction<br>Membrane Protein Function Prediction<br>Functional Domain Impact Prediction |
| Thermal Stability Prediction<br>Fluorescence Intensity Prediction<br>Binding Affinity Prediction<br>Solubility Prediction<br>Expression Level Prediction<br>Enzymatic Activity Prediction<br>Hydrophobicity Prediction<br>Protease Sensitivity Prediction<br>Protein Interaction Strength Prediction<br>Drug Binding Affinity Prediction<br>Protein Stability Change upon Mutation Prediction<br>Protein Ligand Binding Kinetics Prediction<br>Protein Stability in Different Temperatures Prediction<br>Protein Stability in Different Solvents Prediction | **Protein Sequence Design** |
|  | Enzyme Function Optimization<br>Protein Stability Enhancement<br>Protein Folding Prediction<br>Antibody Design<br>Development of Novel Protein Drugs<br>Vaccine Design<br>Protein Engineering<br>Synthetic Antibody Fragments Design |
| **Residue-level Classification** |  |
| Secondary Structure Prediction<br>Binding Site Prediction |  |

**Supplementary Table 2:** Model architectures for Saprot (35M, 650M, and 1.3B) and ESM-2 (35M and 650M) models. Saprot-Seq (35M) was primarily used for ablation studies, employing the same training procedure (including all hyperparameters, network architecture, and training dataset) as Saprot (35M), but with only the AA tokens visible (i.e., masking all 3Di token components).

| Model | Structural information | Parameters | Layers | Embedding size |
| --- | --- | --- | --- | --- |
| Saprot | True | 35M | 12 | 480 |
|  |  | 650M | 33 | 1280 |
|  |  | 1.3B | 66 | 1280 |
| Saprot-Seq | False | 35M | 12 | 480 |
| ESM-2 | False | 35M | 12 | 480 |
|  |  | 650M | 33 | 1280 |

**Supplementary Table 3:** Comparison of Saprot (35M) and ESM-2 (35M) across seven fine-tuning tasks (with protein structures), employing two data splitting strategies: a conventional 30% sequence identity split widely adopted in literature and a 30% LDDT-based structure split introduced by of ProteinShake. To more clearly demonstrate the effectiveness of SA tokens, we also evaluated Saprot-Seq—a variant of Saprot that preserves the exact same architecture and is pre-trained on the same dataset with the same hyperparameters. The only difference is that Saprot-Seq takes only the amino acid sequence as input by masking the structural components (i.e., 3Di information) with “#” symbols. By comparing Saprot and Saprot-Seq, we can clearly see the performance improvement by adding the 3Di structural tokens.

| Split by 30% sequence identity |  |  |  |  |
| --- | --- | --- | --- | --- |
| Task | Metric | ESM-2 (35M) | Saprot-Seq (35M) | Saprot (35M) |
| Thermostability | Spearman’s $\rho$ | 0.669 | 0.672 | **0.692** |
| Metal ion binding | Acc% | 73.08 | 73.23 | **74.29** |
| Subcellular localization | Acc% | 76.58 | 76.67 | **78.09** |
| Binary localization | Acc% | 91.60 | 91.16 | **91.97** |
| Binding site detection | Mcc | 0.609 | 0.507 | **0.627** |
| Structural similarity | Spearman’s $\rho$ | 0.676 | 0.657 | **0.718** |
| Structure class | Acc% | 68.73 | 66.63 | **69.02** |
| Split by 30% structure similarity |  |  |  |  |
| Task | Metric | ESM-2 (35M) | Saprot-Seq (35M) | Saprot (35M) |
| Thermostability | Spearman’s $\rho$ | 0.618 | 0.623 | **0.629** |
| Metal ion binding | Acc% | 68.75 | 65.56 | **72.22** |
| Subcellular localization | Acc% | 76.61 | 76.39 | **77.19** |
| Binary localization | Acc% | 91.47 | 91.71 | **91.82** |
| Binding site detection | Mcc | 0.271 | 0.283 | **0.434** |
| Structural similarity | Spearman’s $\rho$ | 0.719 | 0.707 | **0.754** |
| Structure class | Acc% | 36.55 | 34.46 | **37.55** |

**Supplementary Table 4:** Performance comparison of Saprot with 650M and 1.3B versions. Due to computational and memory constraints, we evaluated the Saprot 650M and 1.3B models on seven randomly selected tasks (three mutation effect prediction tasks and four supervised fine-tuning tasks). For the supervised tasks, we implemented a lightweight fine-tuning approach that only retrains the prediction heads while keeping the backbones frozen. This approach was necessary as the computational requirements of Saprot 1.3B made both standard full fine-tuning and LoRA-based methods difficult, even with 80GB GPU memory available. Additionally, considering the relatively small performance gains observed and computational resource constraints, we did not pursue further comparative analyses on more tasks. These findings suggest that merely scaling up model size in protein language models may encounter diminishing returns, potentially indicating a performance ceiling. Scaling AF2’s structure predictions to billions of proteins could potentially yield larger improvements with a 1.3B parameter model; however, this exploration extends beyond the scope of this paper.

| Task | Metric | Saprot (650M) | Saprot (1.3B) |
| --- | --- | --- | --- |
| ProteinGym | Spearman’s $\rho$ | 0.457 | **0.460** |
| Mega-scale | Spearman’s $\rho$ | 0.574 | **0.588** |
| ClinVar | AUC | 0.909 | **0.910** |
| Thermostability | Spearman’s $\rho$ | 0.659 | **0.660** |
| Subcellular localization | Acc% | 82.89 | **83.11** |
| Stability | Spearman’s $\rho$ | **0.735** | 0.732 |
| Binary localization | Acc% | 87.24 | **89.59** |

**Supplementary Table 5:** Comparison between ProstT5 and Saprot in several tasks. ProstT5 is a seq2seq T5 model where the input, if it is an amino acid sequence, corresponds to the output of the 3Di sequence, and vice versa. In general, Saprot is comparable to ProstT5 in the protein inverse folding task, but substantially outperform it on other tasks, e.g., zero-shot mutational effect prediction and supervised learning tasks.

| Task / Dataset | Metric | ProstT5 (1.2B) | Saprot (650M) |
| --- | --- | --- | --- |
| ClinVar | AUC | 0.620 | **0.909** |
| Mega-scale | Spearman's $\rho$ | 0.194 | **0.574** |
| Subcellular localization | Acc% | 83.26 | **86.18** |
| Binary localization | Acc% | 94.11 | **94.67** |
| Protein inverse folding | pLDDT | 76.24 | **77.96** |
|  | RMSD | 2.28 | **2.07** |
|  | TM-score | 0.78 | **0.82** |
|  | Sequence recovery | 33.72 | **36.15** |

**Supplementary Table 6:** Supervised downstream dataset descriptions after data preprocessing. For **Training with AF2 structure** and **Training with PDB structure**, datasets are partitioned using a 70% LDDT threshold as recommended by the ProteinShake paper. For **Training without structure**, we retain the original splits provided in the official literature. All these datasets are mutation-related variants for a specific protein. Additionally, in the **Training with AF2 structure** setting, proteins lacking corresponding structures in the AlphaFold DB were excluded from both the training and testing sets. Acc% represents classification accuracy, while Mcc denotes the Matthews Correlation Coefficient.

| Dataset | Evaluation Metric | Train | Valid | Test |
| --- | --- | --- | --- | --- |
| **Training with AF2 structure** |  |  |  |  |
| **Thermostability [61]** | Spearman’s $\rho$ | 5310 | 706 | 706 |
| **Subcellular localization [62]** | Acc% | 10414 | 1368 | 1368 |
| **Binary localization [62]** | Acc% | 6707 | 698 | 807 |
| **Training with PDB structure** |  |  |  |  |
| **Structural similarity [31]** | Spearman’s $\rho$ | 300699 | 4559 | 4850 |
| **Structure class [31]** | Acc% | 7990 | 955 | 1005 |
| **Metal ion binding [72]** | Acc% | 5797 | 719 | 719 |
| **Binding site detection [31]** | Mcc | 2368 | 442 | 464 |
| **Training without structure** |  |  |  |  |
| **Fluorescence [30]** | Spearman’s $\rho$ | 20963 | 5235 | 25517 |
| **Stability [30]** | Spearman’s $\rho$ | 53614 | 2512 | 12851 |
| **AAV [61]** | Spearman’s $\rho$ | 22246 | 2462 | 50432 |
| **β-lactamase [32]** | Spearman’s $\rho$ | 4158 | 520 | 520 |

**Supplementary Table 7:**
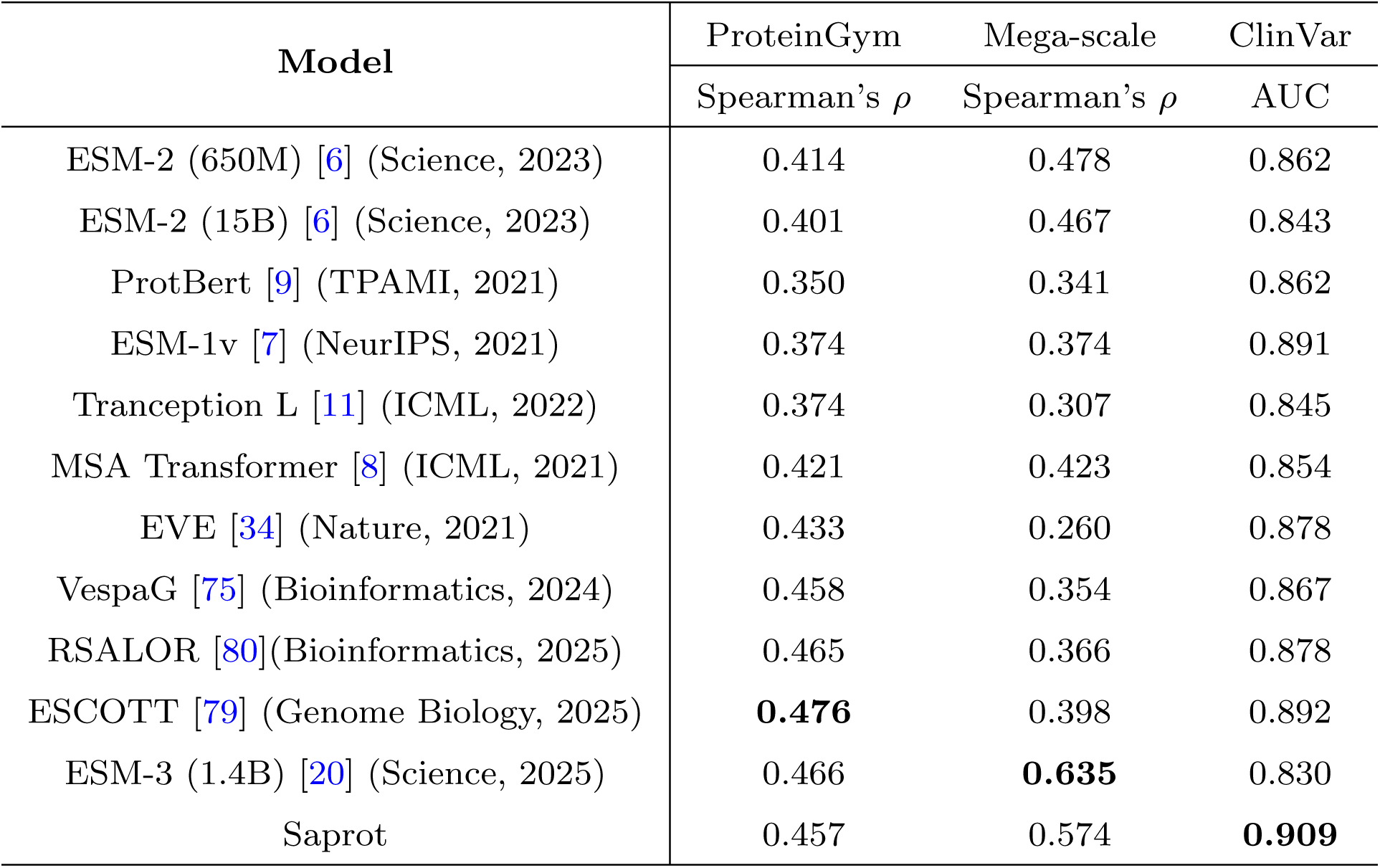
We present additional baseline results for the zero-shot mutational effect prediction task. Notably, VespaG, RSALOR, ESCOTT, and ESM-3 represent recent advances (preprints are published later than Saprot) that have achieved higher rankings than Saprot on the ProteinGym leaderboard. However, their superior performance appears to be dataset-specific, as they fail to outperform Saprot on other mutation effect prediction datasets. In particular, VespaG, RSALOR, and ESCOTT demonstrate substantially inferior performance compared to Saprot on both the Mega-scale and ClinVar datasets. Saprot is comparable to ESM-3, which is expected because both models employ the “amino acid + structural token” paradigm pioneered by Saprot.

**Supplementary Table 8:**
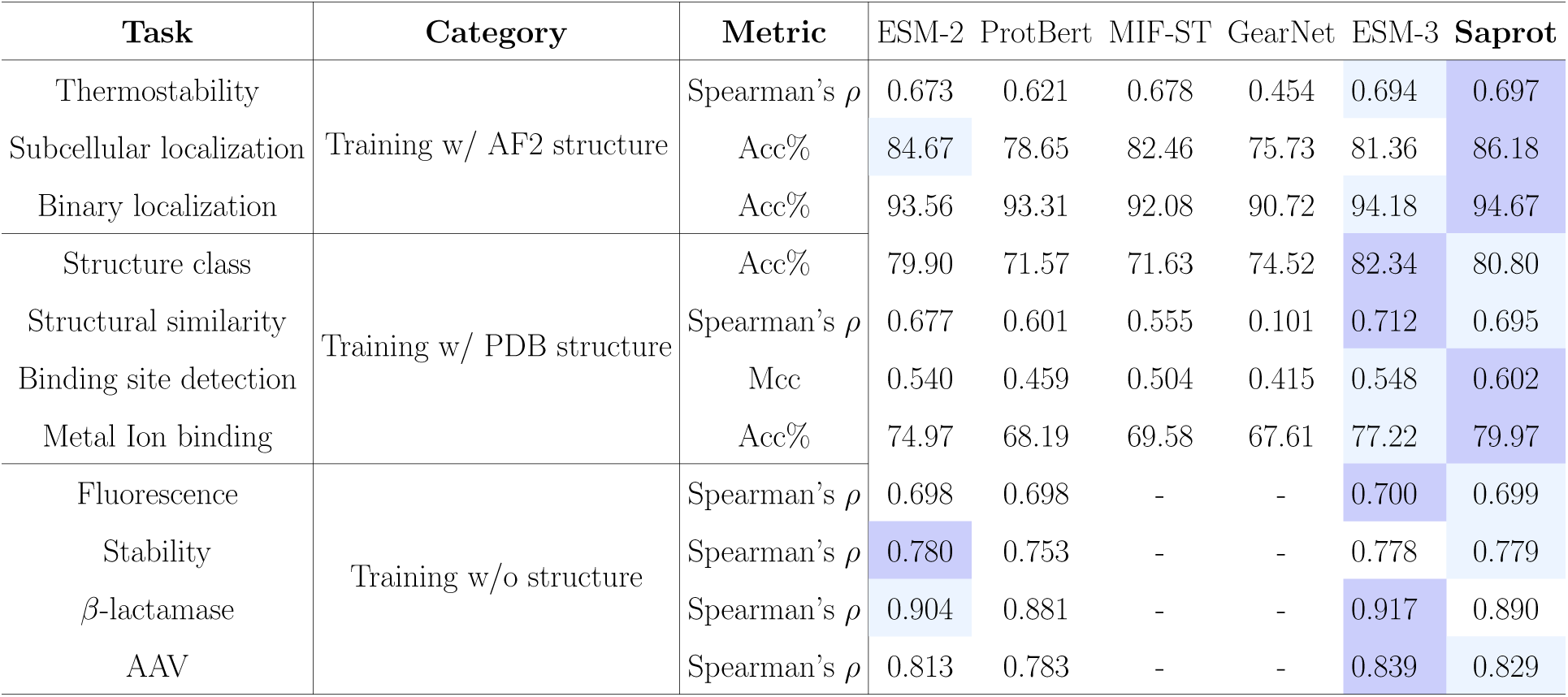
More baseline results on supervised learning tasks. MIF-ST and GearNet are not applicable when protein structures are not available as input. ESM-2 and Saprot have 650M parameters, while ProtBert and ESM-3 have 420M and 1.4B parameters, respectively.

| Task | Category | Metric | ESM-2 | ProtBert | MIF-ST | GearNet | ESM-3 | Saprot |
| --- | --- | --- | --- | --- | --- | --- | --- | --- |
| Thermostability | Training w/ AF2 structure | Spearman's $\rho$ | 0.673 | 0.621 | 0.678 | 0.454 | 0.694 | 0.697 |
| Subcellular localization |  | Acc% | 84.67 | 78.65 | 82.46 | 75.73 | 81.36 | 86.18 |
| Binary localization |  | Acc% | 93.56 | 93.31 | 92.08 | 90.72 | 94.18 | 94.67 |
| Structure class | Training w/ PDB structure | Acc% | 79.90 | 71.57 | 71.63 | 74.52 | 82.34 | 80.80 |
| Structural similarity | | Spearman's $\rho$ | 0.677 | 0.601 | 0.555 | 0.101 | 0.712 | 0.695 |
| Binding site detection |  | Mcc | 0.540 | 0.459 | 0.504 | 0.415 | 0.548 | 0.602 |
| Metal Ion binding |  | Acc% | 74.97 | 68.19 | 69.58 | 67.61 | 77.22 | 79.97 |
| Fluorescence | Training w/o structure | Spearman's $\rho$ | 0.698 | 0.698 | - | - | 0.700 | 0.699 |
| Stability | | Spearman's $\rho$ | 0.780 | 0.753 | - | - | 0.778 | 0.779 |
| β-lactamase | | Spearman's $\rho$ | 0.904 | 0.881 | - | - | 0.917 | 0.890 |
| AAV | | Spearman's $\rho$ | 0.813 | 0.783 | - | - | 0.839 | 0.829 |

**Supplementary Table 9:** Performance evaluation of protein sequence design on 100 randomly selected protein structures from the most recent releases (December 2024). The PDB ids of these structures are available at https://drive.google.com/file/d/1gjZPY63JsA4RbycAfhvBgjdhSYTuG55o/view?usp=sharing.

| Model | pLDDT | RMSD (Å) | TM-score | Sequence recovery (Acc%) |
| --- | --- | --- | --- | --- |
| ProteinMPNN | 82.60 | 1.77 | 0.88 | 48.47 |
| Saprot (AF2 + CATH) | 82.30 | 2.02 | 0.87 | 45.19 |

**Supplementary Table 10:** All data points used to plot Fig. 2e. The training set was evenly divided into multiple parts, with each model trained on a different subset. Training was conducted independently several times, and the average results were computed. ‘Model agg’ refers to the aggregation of predictions generated by individual models (see Methods 1.13).

| Model | Thermostability (Spearman's $\rho$) | | | | | | Subcellular Localization (Acc%) | | | | | |
| --- | --- | --- | --- | --- | --- | --- | --- | --- | --- | --- | --- | --- |
|  | Sample size: 1062 |  |  |  |  |  | Sample size: 2083 |  |  |  |  |  |
|  | Independent result |  |  |  |  | Average result | Independent result |  |  |  |  | Average result |
| Model_0 | 0.617 | 0.643 | 0.589 | 0.616 | 0.632 | 0.619 | 79.97 | 80.26 | 80.34 | 81.80 | 79.46 | 80.37 |
| Model_1 | 0.647 | 0.633 | 0.641 | 0.654 | 0.641 | 0.643 | 79.61 | 80.85 | 80.77 | 79.68 | 82.02 | 80.59 |
| Model_2 | 0.638 | 0.661 | 0.647 | 0.641 | 0.644 | 0.646 | 81.14 | 79.97 | 82.68 | 81.65 | 80.70 | 81.23 |
| Model_3 | 0.636 | 0.667 | 0.648 | 0.649 | 0.651 | 0.650 | 80.34 | 80.77 | 82.09 | 82.38 | 83.92 | 81.90 |
| Model_4 | 0.653 | 0.676 | 0.670 | 0.653 | 0.632 | 0.657 | 82.38 | 82.89 | 84.87 | 83.26 | 82.53 | 83.19 |
| Model_agg | 0.689 | 0.700 | 0.690 | 0.689 | 0.683 | 0.690 | 83.92 | 83.77 | 85.44 | 84.69 | 85.09 | 84.58 |

**Supplementary Table 11:**
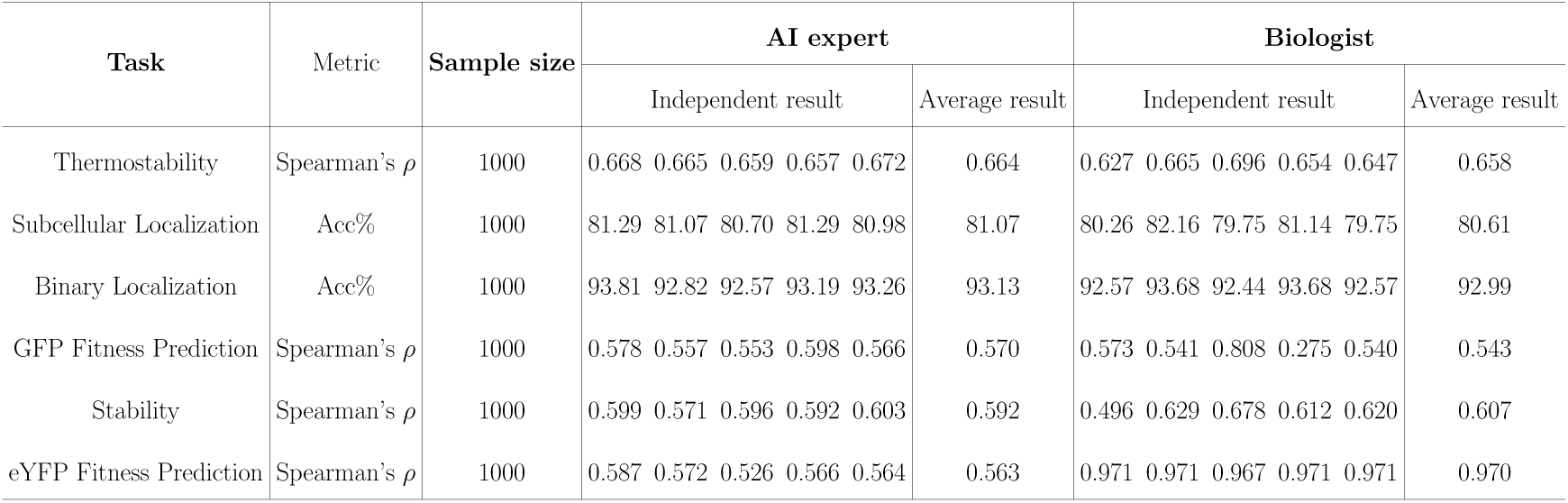
All data points used to plot Fig. 2g. Materials for user study are available at https://drive.google.com/file/d/1LdGRnwt2lttnszNBAJ0F967A8rguPq8b/view?usp=sharing.

**Supplementary Table 12:** The 20 highest-ranked Mth (xylanase) variants predicted by ColabSaprot were expressed in *Pichia pastoris*. Results showed that 13 mutants exhibited increased specific activity compared to the wild-type enzyme. The most effective variant, R59S, reached 2.55-fold of the wild-type (WT) activity. Several other variants, including S225N, G22A, A194F, and F212N, reached approximately 2-fold of the wild-type activity. All of the assays were repeated at least three times. The structure used for prediction is available at https://drive.google.com/file/d/1TgEYZQLwXmshi1X93GAji7k5AD-rZCd/view?usp=sharing.

| Mutation | Predicted score | Specific activity (U/*ml*)<br>(wild type=5.9101) | Enhancement factor |
| --- | --- | --- | --- |
| F212Q | 7.5327 | 4.4419 | 0.7516 |
| R59T | 5.3584 | 0.4921 | 0.0833 |
| F212M | 4.6898 | 4.4720 | 0.7567 |
| F212R | 4.3376 | 3.9598 | 0.6700 |
| N236P | 4.2207 | 8.6526 | 1.4640 |
| S165T | 4.0865 | 7.0325 | 1.1899 |
| A15G | 3.9353 | - | - |
| **R59S** | 3.8820 | **15.1057** | **2.5559** |
| M184V | 3.8267 | 1.0006 | 0.1693 |
| **A194F** | 3.7584 | **11.5114** | **1.9478** |
| **F212N** | 3.7492 | **11.1590** | **1.8881** |
| **S225N** | 3.7452 | **12.9281** | **2.1875** |
| A194L | 3.6962 | 10.4406 | 1.7666 |
| F212L | 3.6802 | 1.1437 | 0.1935 |
| M184T | 3.5455 | 0.4883 | 0.0826 |
| R59Q | 3.4463 | 10.8399 | 1.8341 |
| **G22A** | 3.4332 | **14.6741** | **2.4829** |
| N236G | 3.3842 | 8.0005 | 1.3537 |
| G22V | 3.3259 | 10.1135 | 1.7112 |
| I13P | 3.2729 | - | - |
| F212S | 3.2339 | 8.0726 | 1.3659 |
| D44N | 3.2112 | 7.8896 | 1.3349 |

**Supplementary Table 13:**
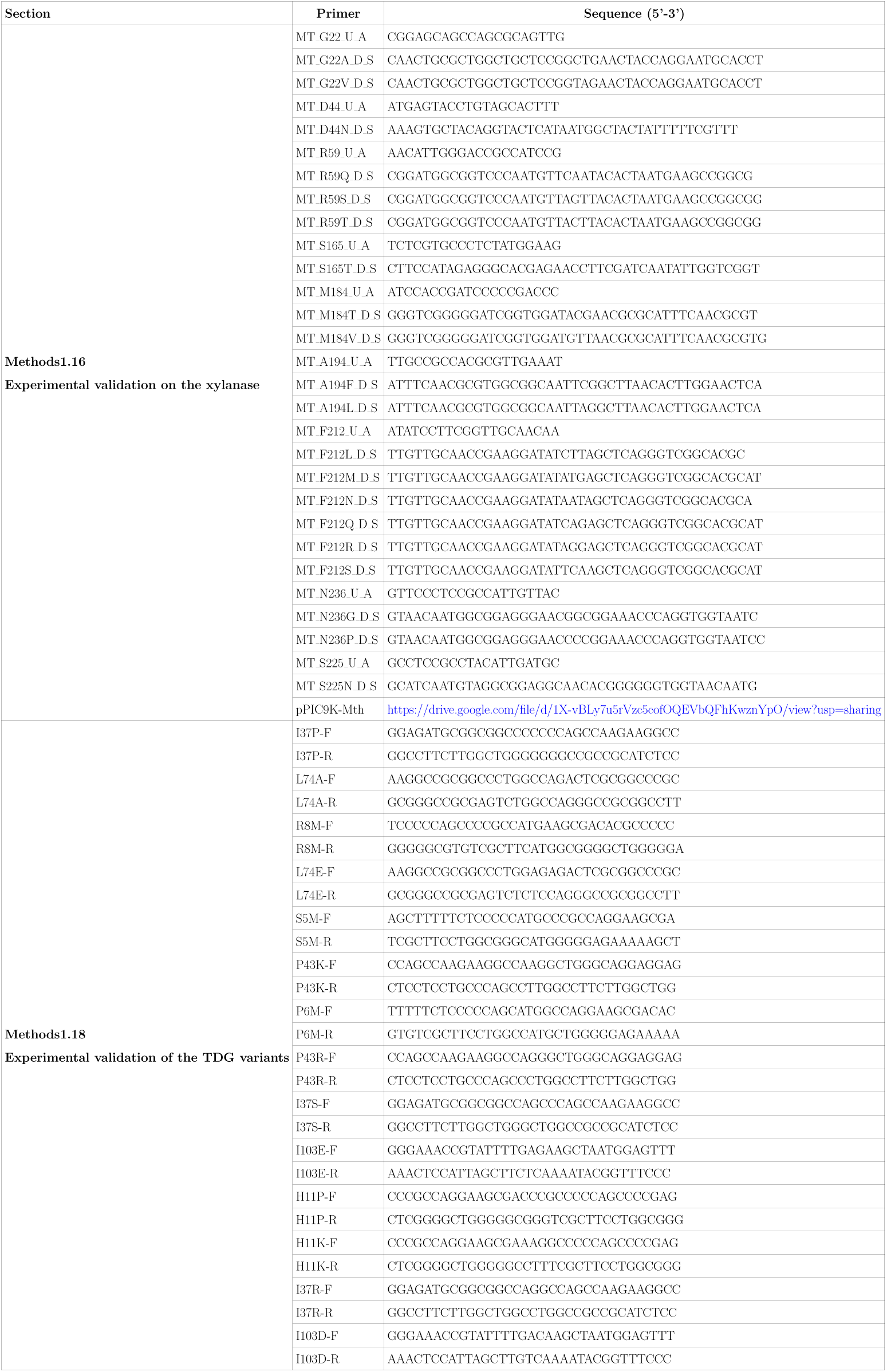
Site-directed mutation primers used in this study.

**Supplementary Table 14:**
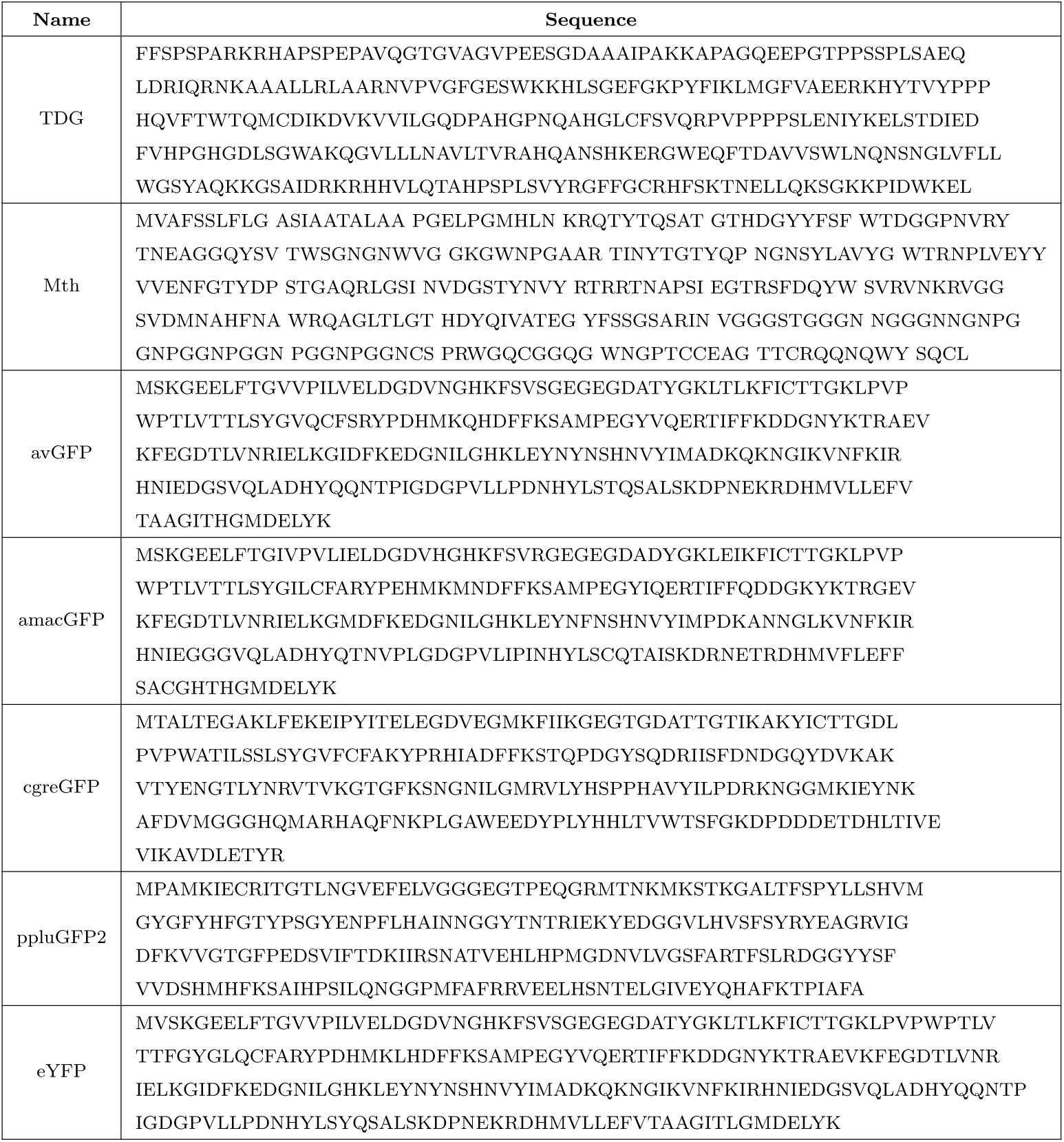
Amino acid sequence of the wild-type protein studied in this experiment.

**Supplementary Table 15:** Validation of the top 20 TDG variants predicted by ColabSaprot. The top 20 TDG variants predicted by ColabSaprot were inserted into the nCas9 protein and co-transfected into HeLa cells along with Dicer-1 sgRNA. Five days after transfection, T*>*G substitutions were analyzed using high-throughput sequencing (HTS). Among the top 20 variants, 17 exhibited higher editing efficiencies compared to the wild-type. Specifically, three variants (L74E, L74Q, and H11K) demonstrated nearly double the editing efficiency relative to the wild-type.

| Mutation | Predicted score by ColabSaprot | T>G%<br>(wildtype=10.3865) | Enhancement factor |
| --- | --- | --- | --- |
| I37P | 4.8993 | 15.7850 | 1.5198 |
| L74A | 4.7470 | 12.9784 | 1.2495 |
| R8M | 3.8026 | 14.7522 | 1.4203 |
| **L74E** | 3.6393 | **20.4948** | **1.9732** |
| S5M | 3.6230 | 15.0282 | 1.4469 |
| P43K | 3.3585 | 14.7687 | 1.4219 |
| P6M | 3.3499 | 15.5822 | 1.5002 |
| P43R | 3.2832 | 13.6750 | 1.3166 |
| I37S | 3.2649 | 16.4798 | 1.5866 |
| I103E | 3.2008 | 11.9370 | 1.1493 |
| H11P | 3.1735 | 14.6748 | 1.4129 |
| **H11K** | 3.1643 | **18.0260** | **1.7355** |
| I37R | 3.1442 | 14.8948 | 1.4341 |
| I103D | 3.1406 | 6.9425 | 0.6684 |
| F1M | 3.0996 | 6.0477 | 0.5823 |
| **L74Q** | 3.0893 | **19.1058** | **1.8395** |
| H11A | 2.9147 | 17.1734 | 1.6534 |
| P4M | 2.7864 | 15.2689 | 1.4701 |
| I37A | 2.7000 | 16.3456 | 1.5737 |
| K252L | 2.6721 | 1.2497 | 0.1203 |

**Supplementary Table 16:**
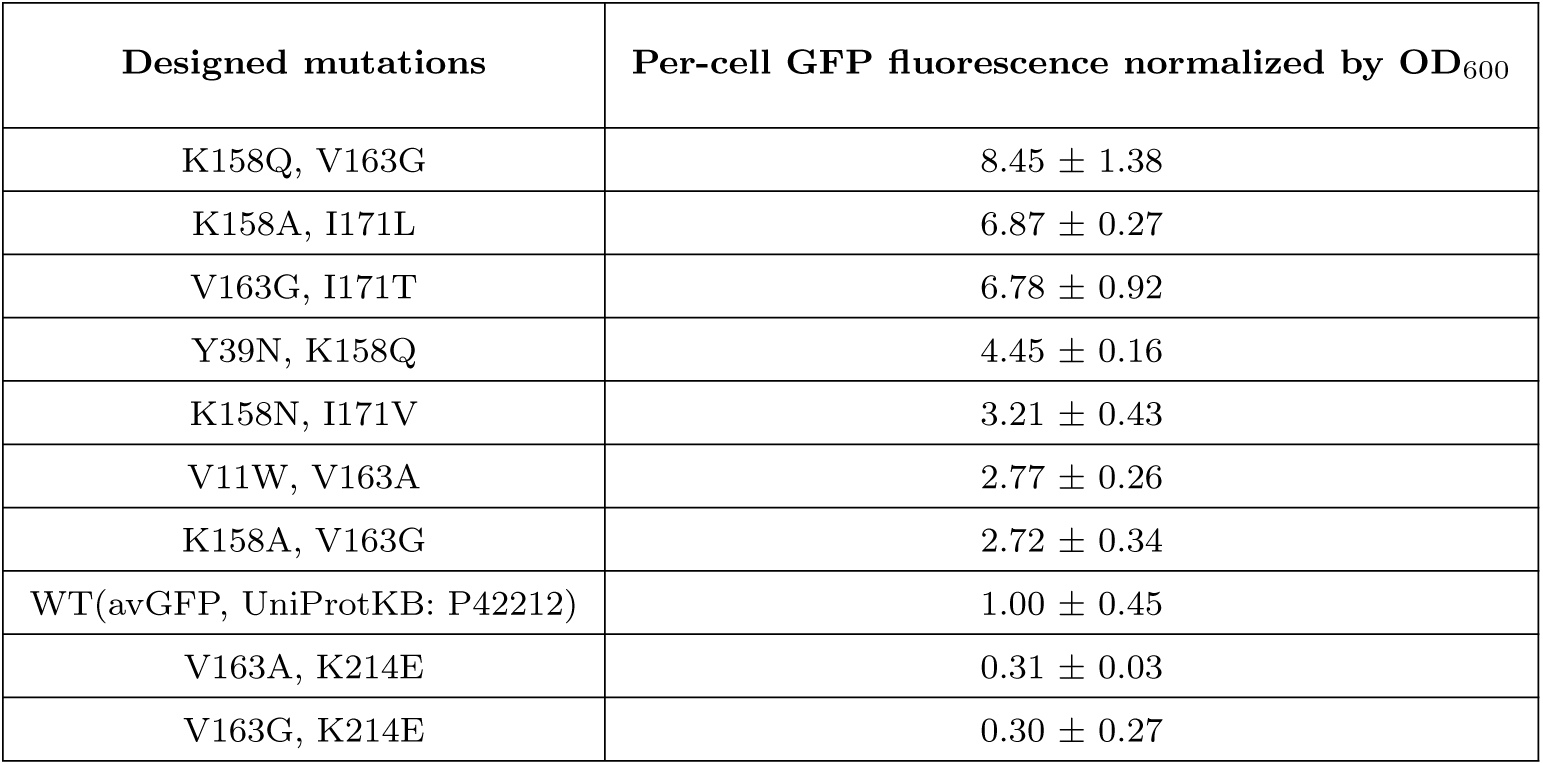
Top 9 avGFP mutants predicted by Saprot and experimental performance. Fluorescence intensities were normalized relative to the wild-type (WT) parent, which was set as 1. The best-performing variant reached over 8-fold of the WT fluorescence intensity.

| Designed mutations | Per-cell GFP fluorescence normalized by OD<sub>600</sub> |
| --- | --- |
| K158Q, V163G | 8.45 ± 1.38 |
| K158A, I171L | 6.87 ± 0.27 |
| V163G, I171T | 6.78 ± 0.92 |
| Y39N, K158Q | 4.45 ± 0.16 |
| K158N, I171V | 3.21 ± 0.43 |
| V11W, V163A | 2.77 ± 0.26 |
| K158A, V163G | 2.72 ± 0.34 |
| WT(avGFP, UniProtKB: P42212) | 1.00 ± 0.45 |
| V163A, K214E | 0.31 ± 0.03 |
| V163G, K214E | 0.30 ± 0.27 |

**Supplementary Fig. 1:**
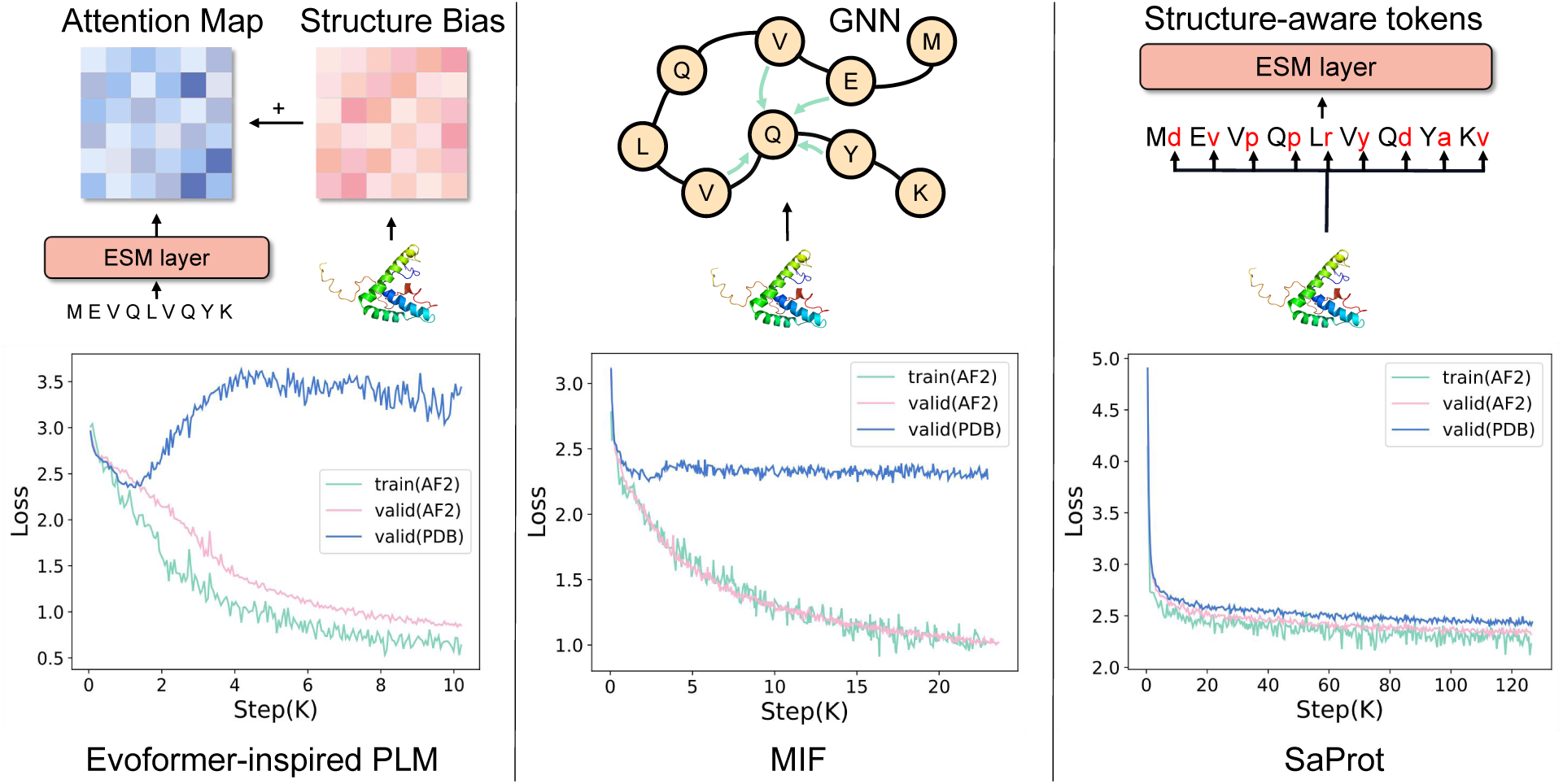
Loss trends for three structure-based PLMs. The training set is the AF2-predicted structures, while in the validation set, one is the AF2 structure and the other is the experimental structure from PDB. The figure is from our Saprot conference version in ICLR 2024.

**Supplementary Fig. 2:**
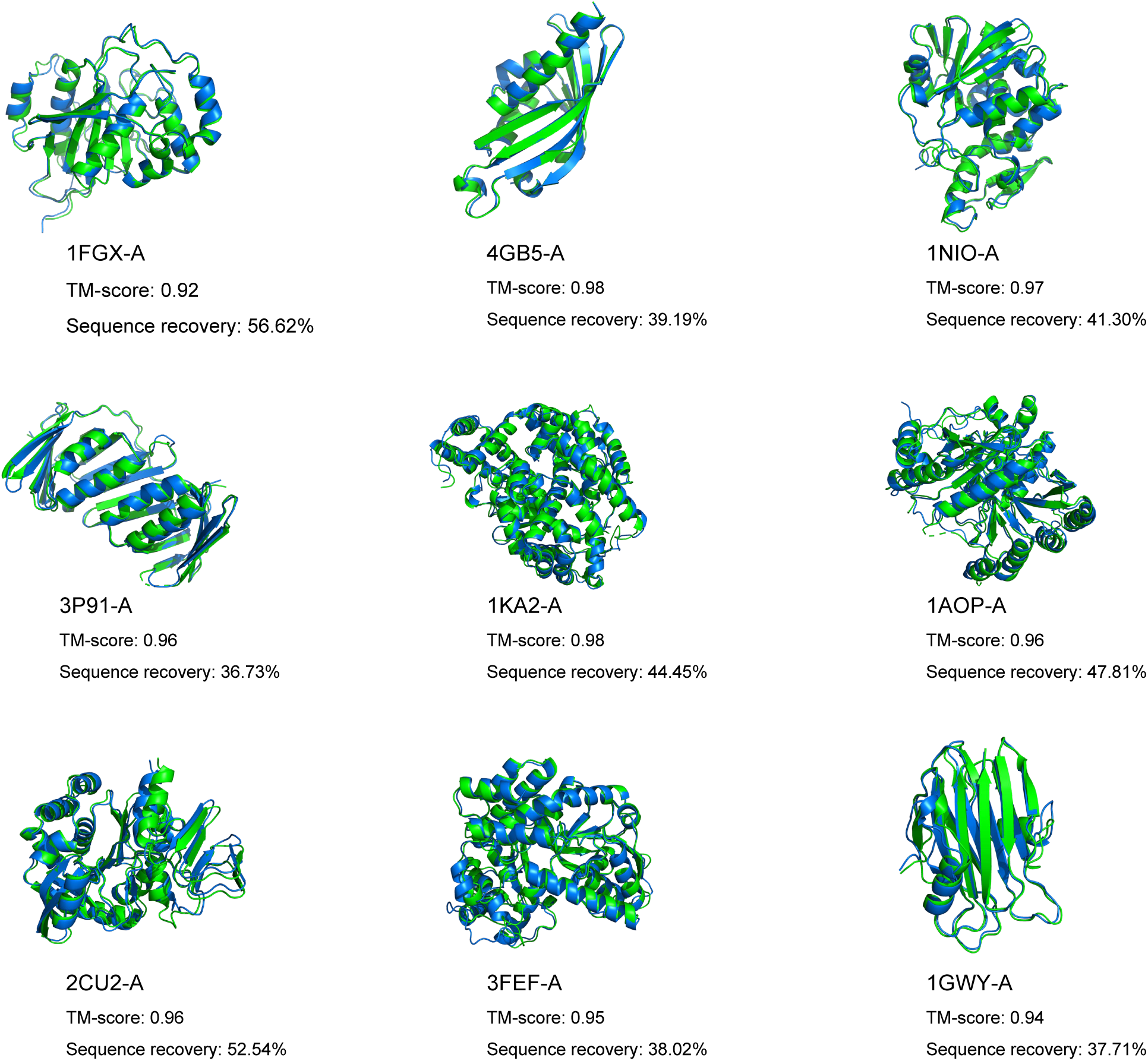
Case studies of protein inverse folding using ColabSaprot (650M). The blue-colored protein structures are predicted using ESMFold, with residue sequences generated by Saprot, and the green-colored structures are the experimentally determined structures. Given the backbone, Saprot is capable of predicting diverse sequences with few residue overlap while maintaining almost the same 3D structure.

**Supplementary Fig. 3:**
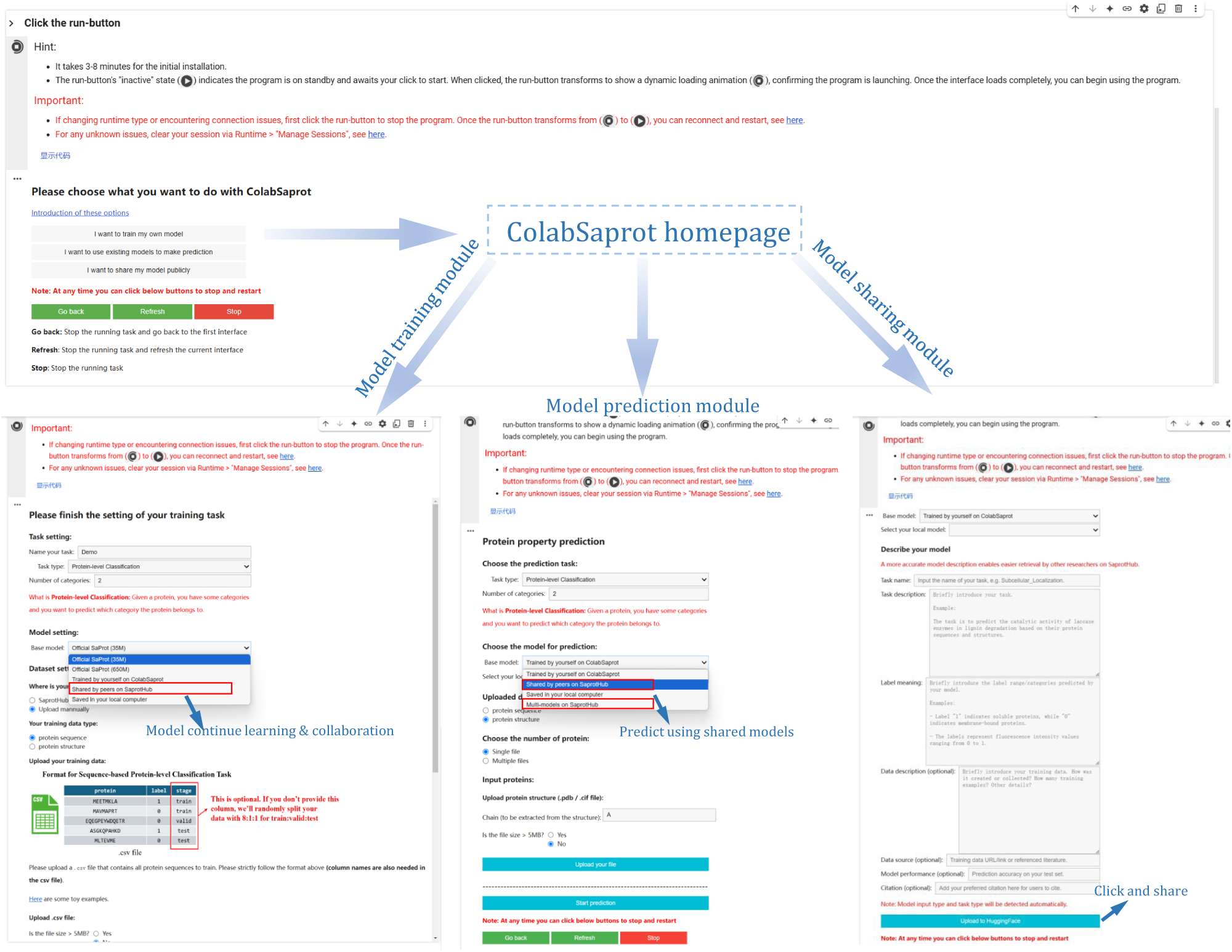
ColabSaprot webpage (https://colab.research.google.com/github/westlake-repl/SaprotHub/blob/main/colab/SaprotHubv2.ipynb?hl=en).

**Supplementary Fig. 4:**
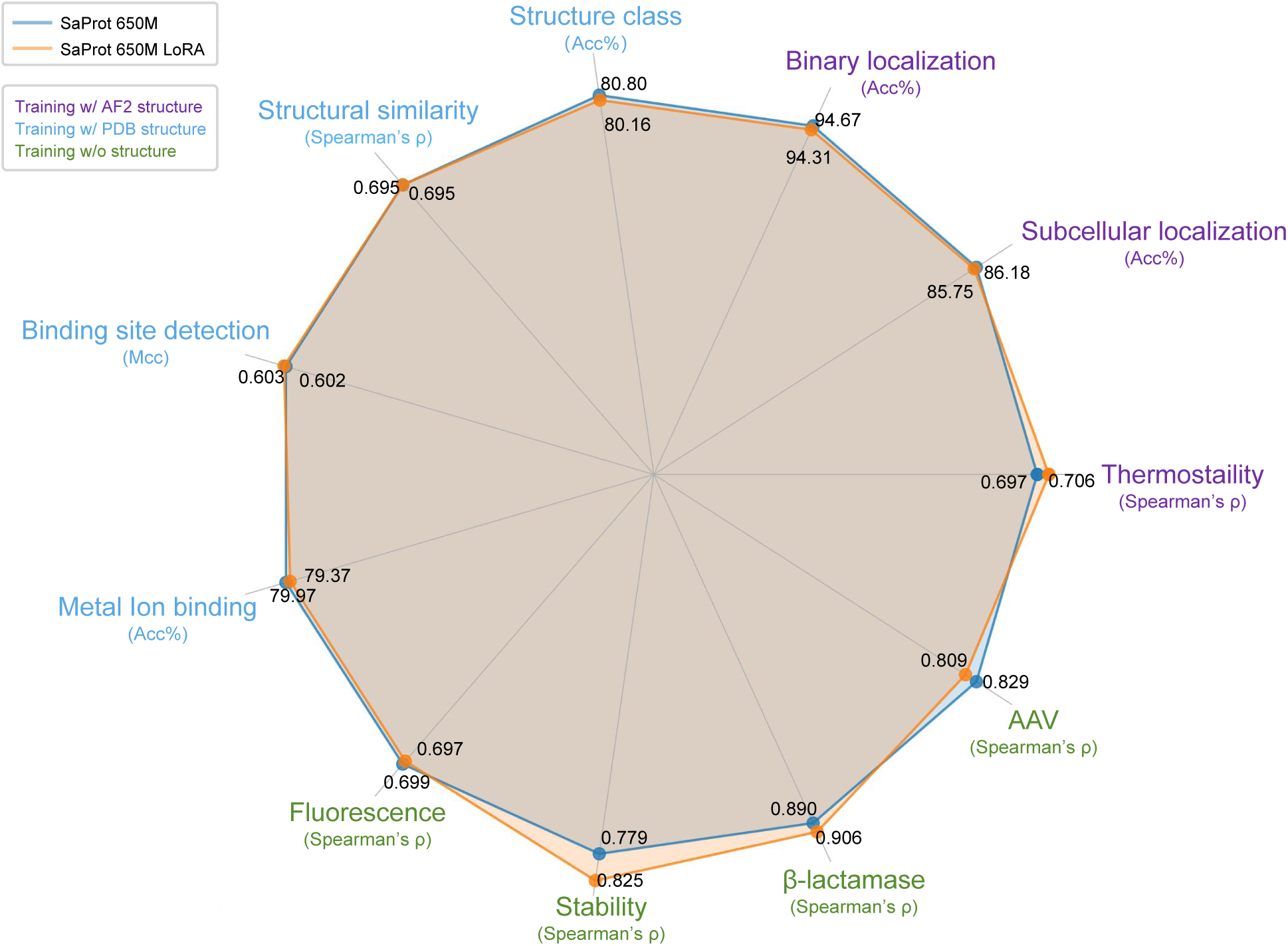
Comparison between full model fine-tuning and parameter efficient fine-tuning. While greatly reducing trainable parameters, fine-tuning Saprot using adapter module achieves competitive performance on all experimental settings, which demonstrates its applicability on various biological tasks.

**Supplementary Fig. 5:**
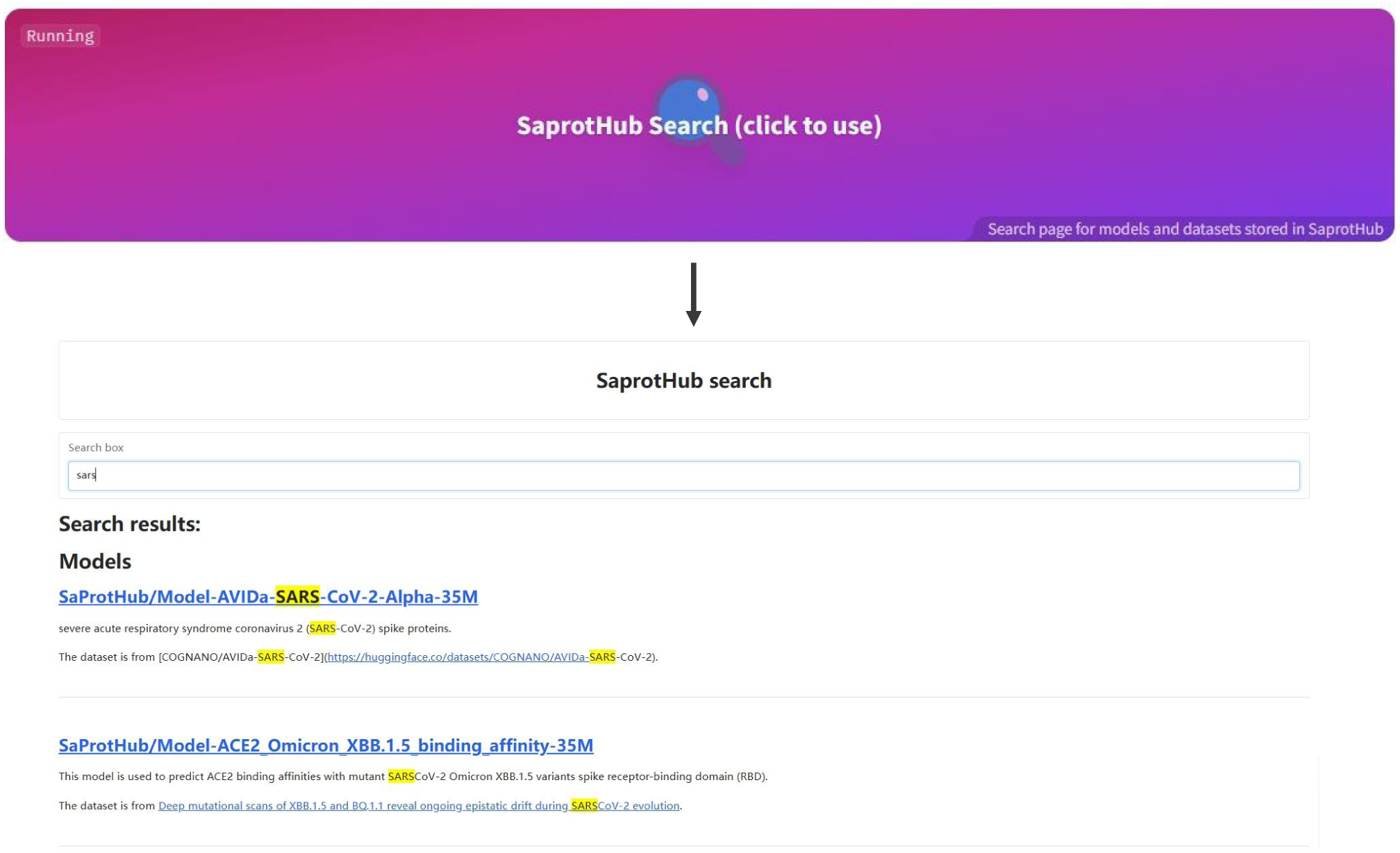
The model search engine (https://huggingface.co/spaces/SaProtHub/SaprotHub-search) on the SaprotHub webpage (https://huggingface.co/SaProtHub).

**Supplementary Fig. 6:**
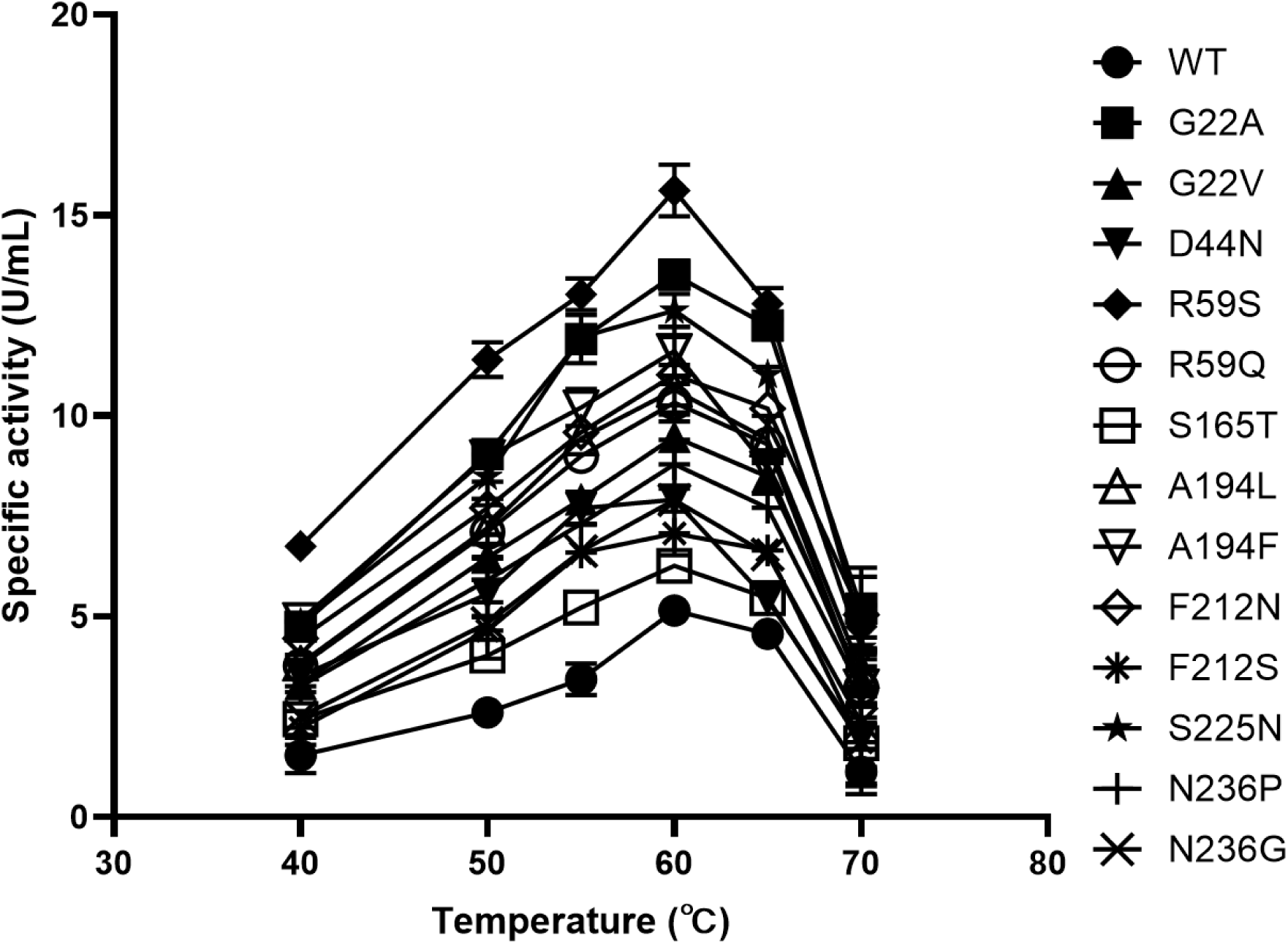
The optimal reaction temperature for both the top-ranked mutants and the wild type (WT) was 60°C. Error bars represent standard deviations, and two replicates were performed.

**Supplementary Fig. 7:**
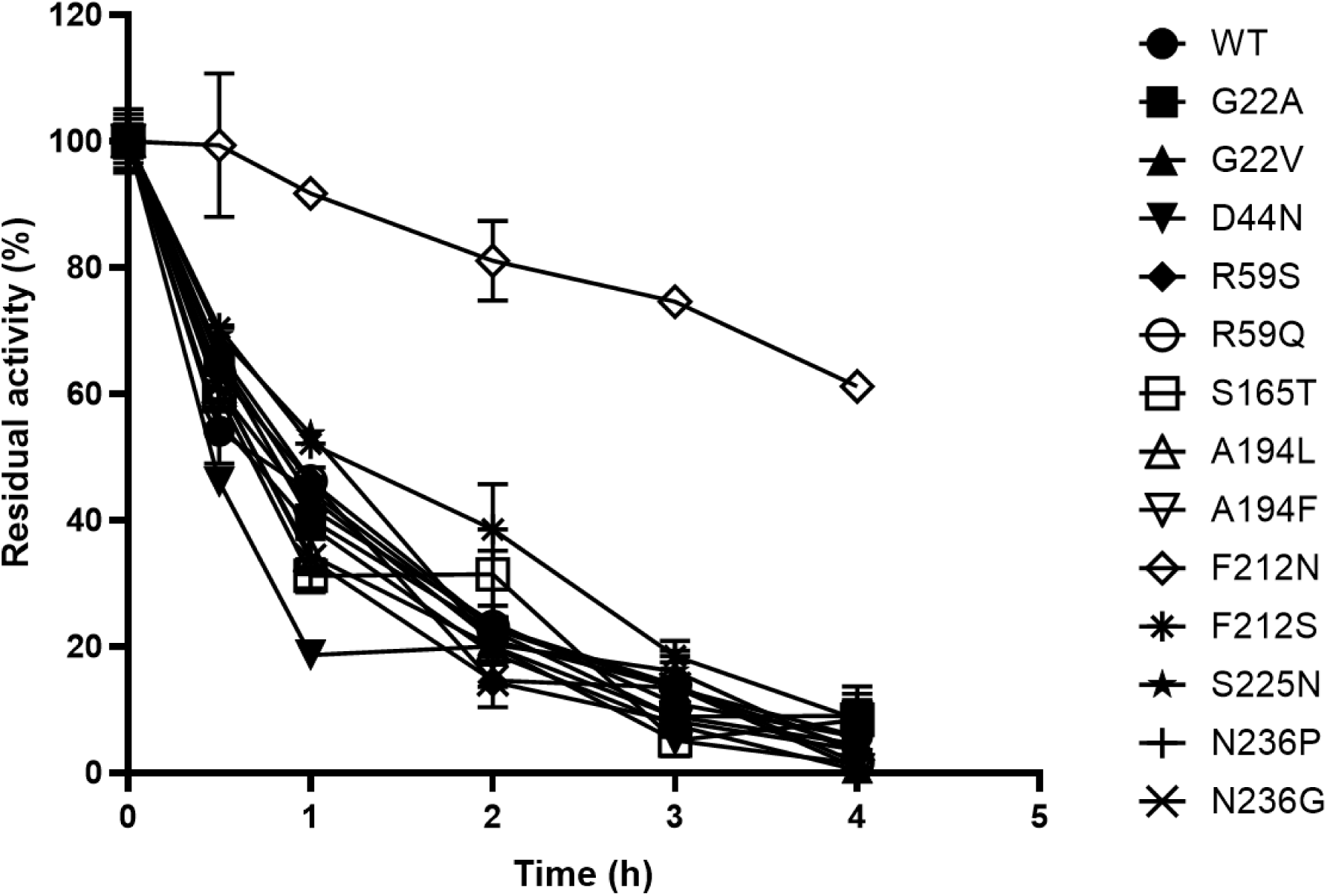
Mutant F212N exhibited a 1.88-fold increase in specific activity compared to the wild-type (WT) enzyme, along with significantly improved thermal stability. After a 4-hour incubation at 60°C, F212N retained over 60% of its initial activity, while the WT enzyme was almost completely inactivated under the same conditions. Error bars represent standard deviations, and two replicates were performed.

## References

[1] Jurgen Drews. Drug discovery: a historical perspective. science, 287(5460):1960–1964, 2000.

[2] François Jacob and Jacques Monod. Genetic regulatory mechanisms in the synthesis of proteins. Journal of molecular biology, 3(3):318–356, 1961.

[3] Michael H Glickman and Aaron Ciechanover. The ubiquitin-proteasome proteolytic pathway: destruction for the sake of construction. Physiological reviews, 2002.

[4] John Jumper, Richard Evans, Alexander Pritzel, Tim Green, Michael Figurnov, Olaf Ronneberger, Kathryn Tunyasuvunakool, Russ Bates, Augustin Žídek, Anna Potapenko, et al. Highly accurate protein structure prediction with alphafold. Nature, 596(7873):583–589, 2021.

[5] Alexander Rives, Joshua Meier, Tom Sercu, Siddharth Goyal, Zeming Lin, Jason Liu, Demi Guo, Myle Ott, C Lawrence Zitnick, Jerry Ma, et al. Biological structure and function emerge from scaling unsupervised learning to 250 million protein sequences. Proceedings of the National Academy of Sciences, 118(15):e2016239118, 2021.

[6] Zeming Lin, Halil Akin, Roshan Rao, Brian Hie, Zhongkai Zhu, Wenting Lu, Nikita Smetanin, Robert Verkuil, Ori Kabeli, Yaniv Shmueli, et al. Evolutionary-scale prediction of atomic-level protein structure with a language model. Science, 379(6637):1123–1130, 2023.

[7] Joshua Meier, Roshan Rao, Robert Verkuil, Jason Liu, Tom Sercu, and Alex Rives. Language models enable zero-shot prediction of the effects of mutations on protein function. Advances in neural information processing systems, 34:29287–29303, 2021.

[8] Roshan M Rao, Jason Liu, Robert Verkuil, Joshua Meier, John Canny, Pieter Abbeel, Tom Sercu, and Alexander Rives. Msa transformer. In International Conference on Machine Learning, pages 8844–8856. PMLR, 2021.

[9] Ahmed Elnaggar, Michael Heinzinger, Christian Dallago, Ghalia Rehawi, Yu Wang, Llion Jones, Tom Gibbs, Tamas Feher, Christoph Angerer, Martin Steinegger, et al. Prottrans: Toward understanding the language of life through self-supervised learning. IEEE transactions on pattern analysis and machine intelligence, 44(10):7112–7127, 2021.

[10] Michael Heinzinger, Konstantin Weissenow, Joaquin Gomez Sanchez, Adrian Henkel, Martin Steinegger, and Burkhard Rost. Prostt5: Bilingual language model for protein sequence and structure. bioRxiv, pages 2023–07, 2023.

[11] Pascal Notin, Mafalda Dias, Jonathan Frazer, Javier Marchena-Hurtado, Aidan N Gomez, Debora Marks, and Yarin Gal. Tranception: protein fitness prediction with autoregressive transformers and inference-time retrieval. In International Conference on Machine Learning, pages 16990–17017. PMLR, 2022.

[12] Erik Nijkamp, Jeffrey A Ruffolo, Eli N Weinstein, Nikhil Naik, and Ali Madani. Progen2: exploring the boundaries of protein language models. Cell systems, 14(11):968–978, 2023.

[13] Noelia Ferruz, Steffen Schmidt, and Birte Hocker. Protgpt2 is a deep unsupervised language model for protein design. Nature communications, 13(1):4348, 2022.

[14] Jin Su, Chenchen Han, Yuyang Zhou, Junjie Shan, Xibin Zhou, and Fajie Yuan. Saprot: Protein language modeling with structure-aware vocabulary. In The Twelfth International Conference on Learning Representations, 2023.

[15] Milot Mirdita, Konstantin Schütze, Yoshitaka Moriwaki, Lim Heo, Sergey Ovchinnikov, and Martin Steinegger. Colabfold: making protein folding accessible to all. Nature methods, 19(6):679–682, 2022.

[16] Edward J Hu, Yelong Shen, Phillip Wallis, Zeyuan Allen-Zhu, Yuanzhi Li, Shean Wang, Lu Wang, and Weizhu Chen. Lora: Low-rank adaptation of large language models. *arXiv preprint arXiv:2106.09685*, 2021.

[17] Jonas Pfeiffer, Andreas Rückĺe, Clifton Poth, Aishwarya Kamath, Ivan Vulíc, Sebastian Ruder, Kyunghyun Cho, and Iryna Gurevych. Adapterhub: A framework for adapting transformers. *arXiv preprint arXiv:2007.07779*, 2020.

[18] James Kirkpatrick, Razvan Pascanu, Neil Rabinowitz, Joel Veness, Guillaume Desjardins, Andrei A Rusu, Kieran Milan, John Quan, Tiago Ramalho, Agnieszka Grabska-Barwinska, et al. Overcoming catastrophic forgetting in neural networks. Proceedings of the national academy of sciences, 114(13):3521–3526, 2017.

[19] Michel van Kempen, Stephanie S Kim, Charlotte Tumescheit, Milot Mirdita, Cameron LM Gilchrist, Johannes Soding, and Martin Steinegger. Foldseek: fast and accurate protein structure search. Biorxiv, pages 2022–02, 2022.

[20] Thomas Hayes, Roshan Rao, Halil Akin, Nicholas James Sofroniew, Deniz Oktay, Zeming Lin, Robert Verkuil, Vincent Quy Tran, Jonathan Deaton, Marius Wiggert, Rohil Badkundri, Irhum Shafkat, Jun Gong, Alexander Derry, Raul Santiago Molina, Neil Thomas, Yousuf Khan, Chetan Mishra, Carolyn Kim, Liam J. Bartie, Patrick D. Hsu, Tom Sercu, Salvatore Candido, and Alexander Rives. Simulating 500 million years of evolution with a language model. bioRxiv, 2024.

[21] Mingchen Li, Yang Tan, Xinzhu Ma, Bozitao Zhong, Huiqun Yu, Ziyi Zhou, Wanli Ouyang, Bingxin Zhou, Liang Hong, and Pan Tan. Prosst: Protein language modeling with quantized structure and disentangled attention. bioRxiv, pages 2024–04, 2024.

[22] Xinyou Wang, Zaixiang Zheng, Fei Ye, Dongyu Xue, Shujian Huang, and Quanquan Gu. Dplm-2: A multimodal diffusion protein language model. *arXiv preprint arXiv:2410.13782*, 2024.

[23] Yang Tan, Ruilin Wang, Banghao Wu, Liang Hong, and Bingxin Zhou. Retrieval-enhanced mutation mastery: Augmenting zero-shot prediction of protein language model. arXiv preprint arXiv:2410.21127, 2024.

[24] Mahdi Pourmirzaei, Farzaneh Esmaili, Mohammadreza Pourmirzaei, Duolin Wang, and Dong Xu. Prot2token: A multi-task framework for protein language processing using autoregressive language modeling. bioRxiv, pages 2024–05, 2024.

[25] Kaiyuan Gao, Yusong Wang, Haoxiang Guan, Zun Wang, Qizhi Pei, John E Hopcroft, Kun He, and Lijun Wu. Tokenizing 3d molecule structure with quantized spherical coordinates. *arXiv preprint arXiv:2412.01564*, 2024.

[26] Xiaohan Lin, Zhenyu Chen, Yanheng Li, Zicheng Ma, Chuanliu Fan, Ziqiang Cao, Shihao Feng, Yi Qin Gao, and Jun Zhang. Tokenizing foldable protein structures with machine-learned artificial amino-acid vocabulary. bioRxiv, pages 2023–11, 2023.

[27] Nikita V Ivanisenko, Tatiana I Shashkova, Andrey Shevtsov, Maria Sindeeva, Dmitriy Umerenkov, and Olga Kardymon. Sema 2.0: web-platform for b-cell conformational epitopes prediction using artificial intelligence. Nucleic Acids Research, page gkae386, 2024.

[28] Jacob Devlin, Ming-Wei Chang, Kenton Lee, and Kristina Toutanova. Bert: Pre-training of deep bidirectional transformers for language understanding. arXiv preprint arXiv:1810.04805, 2018.

[29] Mihaly Varadi, Stephen Anyango, Mandar Deshpande, Sreenath Nair, Cindy Natassia, Galabina Yordanova, David Yuan, Oana Stroe, Gemma Wood, Agata Laydon, et al. Alphafold protein structure database: massively expanding the structural coverage of protein-sequence space with high-accuracy models. Nucleic acids research, 50(D1):D439–D444, 2022.

[30] Roshan Rao, Nicholas Bhattacharya, Neil Thomas, Yan Duan, Peter Chen, John Canny, Pieter Abbeel, and Yun Song. Evaluating protein transfer learning with tape. Advances in neural information processing systems, 32, 2019.

[31] Tim Kucera, Carlos Oliver, Dexiong Chen, and Karsten Borgwardt. Proteinshake: Building datasets and benchmarks for deep learning on protein structures. Advances in Neural Information Processing Systems, 36, 2024.

[32] Minghao Xu, Zuobai Zhang, Jiarui Lu, Zhaocheng Zhu, Yangtian Zhang, Ma Chang, Runcheng Liu, and Jian Tang. Peer: a comprehensive and multi-task benchmark for protein sequence understanding. Advances in Neural Information Processing Systems, 35:35156–35173, 2022.

[33] Brian Hie, Ellen D Zhong, Bonnie Berger, and Bryan Bryson. Learning the language of viral evolution and escape. Science, 371(6526):284–288, 2021.

[34] Jonathan Frazer, Pascal Notin, Mafalda Dias, Aidan Gomez, Joseph K Min, Kelly Brock, Yarin Gal, and Debora S Marks. Disease variant prediction with deep generative models of evolutionary data. Nature, 599(7883):91–95, 2021.

[35] Justas Dauparas, Ivan Anishchenko, Nathaniel Bennett, Hua Bai, Robert J Ragotte, Lukas F Milles, Basile IM Wicky, Alexis Courbet, Rob J de Haas, Neville Bethel, et al. Robust deep learning–based protein sequence design using proteinmpnn. Science, 378(6615):49–56, 2022.

[36] Kotaro Tsuboyama, Justas Dauparas, Jonathan Chen, Elodie Laine, Yasser Mohseni Behbahani, Jonathan J Weinstein, Niall M Mangan, Sergey Ovchinnikov, and Gabriel J Rocklin. Mega-scale experimental analysis of protein folding stability in biology and design. Nature, 620(7973):434–444, 2023.

[37] Pascal Notin, Aaron Kollasch, Daniel Ritter, Lood Van Niekerk, Steffanie Paul, Han Spinner, Nathan Rollins, Ada Shaw, Rose Orenbuch, Ruben Weitzman, et al. Proteingym: large-scale benchmarks for protein fitness prediction and design. Advances in Neural Information Processing Systems, 36, 2024.

[38] Melissa J Landrum, Jennifer M Lee, Mark Benson, Garth R Brown, Chen Chao, Shanmuga Chitipiralla, Baoshan Gu, Jennifer Hart, Douglas Hoffman, Wonhee Jang, et al. Clinvar: improving access to variant interpretations and supporting evidence. Nucleic acids research, 46(D1):D1062–D1067, 2018.

[39] Yang Tan, Wenrui Gou, Bozitao Zhong, Liang Hong, Huiqun Yu, and Bingxin Zhou. Venusx: Unlocking fine-grained functional understanding of proteins. arXiv preprint arXi:2505.11812, 2025.

[40] Shuo Yan, Yuliang Yan, Bin Ma, Chenao Li, Haochun Tang, Jiahua Lu, Minhua Lin, Yuyuan Feng, Hui Xiong, and Enyan Dai. Protap: A benchmark for protein modeling on realistic downstream applications. arXiv preprint arXiv:2506.02052, 2025.

[41] Ziyi Zhou, Liang Zhang, Yuanxi Yu, Banghao Wu, Mingchen Li, Liang Hong, and Pan Tan. Enhancing efficiency of protein language models with minimal wet-lab data through few-shot learning. Nature Communications, 15(1):5566, 2024.

[42] Fengyuan Dai, Yuliang Fan, Jin Su, Chentong Wang, Chenchen Han, Xibin Zhou, Jianming Liu, Hui Qian, Shunzhi Wang, Anping Zeng, et al. Toward de novo protein design from natural language. bioRxiv, pages 2024–08, 2024.

[43] Viacheslav Meshchaninov, Pavel Strashnov, Andrey Shevtsov, Fedor Nikolaev, Nikita Ivanisenko, Olga Kardymon, and Dmitry Vetrov. Diffusion on language model encodings for protein sequence generation.

[44] Tatsuya Sagawa, Eisuke Kanao, Kosuke Ogata, Koshi Imami, and Yasushi Ishihama. Prediction of protein half-lives from amino acid sequences by protein language models. bioRxiv, 2024.

[45] Anton Bushuiev, Roman Bushuiev, Nikola Zadorozhny, Raman Samusevich, Hannes Stark, Jiri Sedlar, Tomáš Pluskal, and Josef Sivic. Training on test proteins improves fitness, structure, and function prediction, 2024.

[46] Xiang Zhuang, Keyan Ding, Tianwen Lyu, Yinuo Jiang, Xiaotong Li, Zhuoyi Xiang, Zeyuan Wang, Ming Qin, Kehua Feng, Jike Wang, et al. Instructbiomol: Advancing biomolecule understanding and design following human instructions. arXiv preprint arXiv:2410.07919, 2024.

[47] Xibin Zhou, Chenchen Han, Yingqi Zhang, Jin Su, Kai Zhuang, Shiyu Jiang, Zichen Yuan, Wei Zheng, Fengyuan Dai, Yuyang Zhou, et al. Decoding the molecular language of proteins with evolla. bioRxiv, pages 2025–01, 2025.

[48] Lei Wang, Xuchao Zhang, Yan Wang, and Zhidong Xue. Ssalign: Ultrafast and sensitive protein structure search at scale. 2025.

[49] Zhaohan Meng, Zaiqiao Meng, and Iadh Ounis. Fusiondti: Fine-grained binding discovery with token-level fusion for drug-target interaction. arXiv preprint arXiv:2406.01651, 2024.

[50] Andrew T McNutt, Abhinav K Adduri, Caleb N Ellington, Monica T Dayao, Eric P Xing, Hosein Mohimani, and David R Koes. Scaling structure aware virtual screening to billions of molecules with sprint. ArXiv, pages arXiv–2411, 2025.

[51] Yan He, Xibin Zhou, Chong Chang, Ge Chen, Weikuan Liu, Geng Li, Xiaoqi Fan, Mingsun Sun, Chensi Miao, Qianyue Huang, et al. Protein language models-assisted optimization of a uracil-n-glycosylase variant enables programmable t-to-g and t-to-c base editing. Molecular Cell.

[52] Adam J Riesselman, John B Ingraham, and Debora S Marks. Deep generative models of genetic variation capture the effects of mutations. Nature methods, 15(10):816–822, 2018.

53. Chloe Hsu, Robert Verkuil, Jason Liu, Zeming Lin, Brian Hie, Tom Sercu, Adam Lerer, and Alexander Rives. Learning inverse folding from millions of predicted structures. In International conference on machine learning, pages 8946–8970. PMLR, 2022.

[54] Samuel Sledzieski, Meghana Kshirsagar, Minkyung Baek, Bonnie Berger, Rahul Dodhia, and Juan Lavista Ferres. Democratizing protein language models with parameter-efficient fine-tuning. bioRxiv, 2023.

[55] Shuai Zeng, Duolin Wang, and Dong Xu. Peft-sp: Parameter-efficient fine-tuning on large protein language models improves signal peptide prediction. bioRxiv, pages 2023–11, 2023.

[56] Samuel Sledzieski, Meghana Kshirsagar, Bonnie Berger, Rahul Dodhia, and Juan Lavista Ferres. Parameter-efficient fine-tuning of protein language models improves prediction of protein-protein interactions.

[57] Duolin Wang, Mahdi Pourmirzaei, Usman L Abbas, Shuai Zeng, Negin Manshour, Farzaneh Esmaili, Biplab Poudel, Yuexu Jiang, Qing Shao, Jin Chen, et al. S-plm: Structure-aware protein language model via contrastive learning between sequence and structure. Advanced Science, 12(5):2404212, 2025.

[58] Jin Su, Xibin Zhou, Xuting Zhang, and Fajie Yuan. Protrek: Navigating the protein universe through tri-modal contrastive learning. bioRxiv, pages 2024–05, 2024.

[59] Ilya Loshchilov and Frank Hutter. Fixing weight decay regularization in adam. CoRR, abs/1711.05101, 2017.

[60] Zuobai Zhang, Minghao Xu, Arian Jamasb, Vijil Chenthamarakshan, Aurelie Lozano, Payel Das, and Jian Tang. Protein representation learning by geometric structure pretraining. arXiv preprint arXiv:2203.06125, 2022.

[61] Christian Dallago, Jody Mou, Kadina E Johnston, Bruce J Wittmann, Nicholas Bhattacharya, Samuel Goldman, Ali Madani, and Kevin K Yang. Flip: Benchmark tasks in fitness landscape inference for proteins. bioRxiv, pages 2021–11, 2021.

[62] Jose Juan Almagro Armenteros, Casper Kaae Sønderby, Søren Kaae Sønderby, Henrik Nielsen, and Ole Winther. Deeploc: prediction of protein subcellular localization using deep learning. Bioinformatics, 33(21):3387–3395, 2017.

[63] Vladimir Gligorijevíc, P Douglas Renfrew, Tomasz Kosciolek, Julia Koehler Leman, Daniel Berenberg, Tommi Vatanen, Chris Chandler, Bryn C Taylor, Ian M Fisk, Hera Vlamakis, et al. Structure-based protein function prediction using graph convolutional networks. Nature communications, 12(1):3168, 2021.

[64] Yan He, Xibin Zhou, Fajie Yuan, and Xing Chang. Protocol to use protein language models predicting and following experimental validation of function-enhancing variants of thymine-n-glycosylase. STAR Protocols, 5(3):103188, 2024.

[65] Mingyang Hu, Fajie Yuan, Kevin Yang, Fusong Ju, Jin Su, Hui Wang, Fei Yang, and Qiuyang Ding. Exploring evolution-aware & -free protein language models as protein function predictors. In S. Koyejo, S. Mohamed, A. Agarwal, D. Belgrave, K. Cho, and A. Oh, editors, Advances in Neural Information Processing Systems, volume 35, pages 38873–38884. Curran Associates, Inc., 2022.

## References

[66] Dexiong Chen, Philip Hartout, Paolo Pellizzoni, Carlos Oliver, and Karsten Borgwardt. Endowing protein language models with structural knowledge. arXiv preprint arXiv:2401.14819, 2024.

[67] Junchen Fu, Fajie Yuan, Yu Song, Zheng Yuan, Mingyue Cheng, Shenghui Cheng, Jiaqi Zhang, Jie Wang, and Yunzhu Pan. Exploring adapter-based transfer learning for recommender systems: Empirical studies and practical insights. In Proceedings of the 17th ACM International Conference on Web Search and Data Mining, pages 208–217, 2024.

[68] Lihao Fu, Yuan Gao, Yongcan Chen, Yanjing Wang, Xiaoting Fang, Shujun Tian, Hao Dong, Yijian Zhang, Zichuan Chen, Zechen Wang, et al. Critical assessment of protein engineering (cape): A student challenge on the cloud. ACS Synthetic Biology, 2024.

[69] Linyuan Gong, Di He, Zhuohan Li, Tao Qin, Liwei Wang, and Tieyan Liu. Efficient training of bert by progressively stacking. In *International conference on machine learning*, pages 2337–2346. PMLR, 2019.

[70] Pedro Hermosilla and Timo Ropinski. Contrastive representation learning for 3d protein structures, 2022.

[71] Neil Houlsby, Andrei Giurgiu, Stanislaw Jastrzebski, Bruna Morrone, Quentin De Laroussilhe, Andrea Gesmundo, Mona Attariyan, and Sylvain Gelly. Parameter-efficient transfer learning for nlp. In International conference on machine learning, pages 2790–2799. PMLR, 2019.

[72] Mingyang Hu, Fajie Yuan, Kevin Yang, Fusong Ju, Jin Su, Hui Wang, Fei Yang, and Qiuyang Ding. Exploring evolution-aware & -free protein language models as protein function predictors. In S. Koyejo, S. Mohamed, A. Agarwal, D. Belgrave, K. Cho, and A. Oh, editors, Advances in Neural Information Processing Systems, volume 35, pages 38873–38884. Curran Associates, Inc., 2022.

[73] John Ingraham, Vikas Garg, Regina Barzilay, and Tommi Jaakkola. Generative models for graph-based protein design. Advances in neural information processing systems, 32, 2019.

[74] Rabeeh Karimi Mahabadi, James Henderson, and Sebastian Ruder. Compacter: Efficient low-rank hypercomplex adapter layers. Advances in Neural Information Processing Systems, 34:1022–1035, 2021.

[75] Céline Marquet, Julius Schlensok, Marina Abakarova, Burkhard Rost, and Elodie Laine. Expert-guided protein language models enable accurate and blazingly fast fitness prediction. Bioinformatics, 40(11):btae621, 2024.

[76] Christine A Orengo, Alex D Michie, Susan Jones, David T Jones, Mark B Swindells, and Janet M Thornton. Cath–a hierarchic classification of protein domain structures. Structure, 5(8):1093–1109, 1997.

[77] Robert Schmirler, Michael Heinzinger, and Burkhard Rost. Fine-tuning protein language models boosts predictions across diverse tasks. bioRxiv, pages 2023–12, 2023.

[78] Amelie Schreiber. Esmbind and qbind: Lora, qlora, and esm-2 for predicting binding sites and post translational modification. bioRxiv, pages 2023–11, 2023.

[79] Mustafa Tekpinar, et al. Prescott: a population aware, epistatic and structural model accurately 575 predicts missense effect. medrxiv. 2024.

[80] Matsvei Tsishyn, Pauline Hermans, Marianne Rooman, and Fabrizio Pucci. Residue conservation and solvent accessibility are (almost) all you need for predicting mutational effects in proteins. Bioinformatics, 41(6):btaf322, 2025.

[81] Aaron Van Den Oord, Oriol Vinyals, et al. Neural discrete representation learning. Advances in neural information processing systems, 30, 2017.

[82] Kevin K Yang, Niccolo Zanichelli, and Hugh Yeh. Masked inverse folding with sequence transfer for protein representation learning. Protein Engineering, Design and Selection, 36:gzad015, 2023.

[83] Fajie Yuan, Xiangnan He, Alexandros Karatzoglou, and Liguang Zhang. Parameter-efficient transfer from sequential behaviors for user modeling and recommendation. In Proceedings of the 43rd International ACM SIGIR conference on research and development in Information Retrieval, pages 1469–1478, 2020.

[84] Gengmo Zhou, Zhifeng Gao, Qiankun Ding, Hang Zheng, Hongteng Xu, Zhewei Wei, Linfeng Zhang, and Guolin Ke. Uni-mol: A universal 3d molecular representation learning framework. 2023.

